# Genic and phylogenomic discordances reveal conflicting hybridization episodes in temperate Loliinae grasses

**DOI:** 10.1101/2024.10.25.620283

**Authors:** María Fernanda Moreno-Aguilar, Juan Viruel, Diana Calderón, Itziar Arnelas, Aminael Sánchez-Rodríguez, Wenli Chen, Juan Camilo Ospina, Gloria Martínez-Sagarra, Juan Antonio Devesa, Alan Stewart, Pilar Catalán

## Abstract

The grass subtribe Loliinae has great ecological and economic importance, as it includes community-dominant species of world mountain grasslands and the most extensively cultivated pasture, fodder and lawn grasses (fescues, ryegrasses). Resolving the phylogeny of recently evolved Loliinae lineages has proven challenging due to frequent introgressions and polyploidizations that occurred throughout their history. Evolutionary reconstruction based on genome-wide data sets is useful for large groups of species, enabling recovering phylogenetic trees, contrasting their topologies, unraveling the origins of the discordances, and testing hypotheses on their reticulate history. Here we present the first target-capture phylogeny of Loliinae using 234 single-copy nuclear coding loci for a large sample of 132 representative taxa, covering all its 29 evolutionary lineages. Additionally, we have completed organelle (plastome) sequences to elucidate the hybridogenic speciation history of the Loliinae from complementary genome sources. Concatenated maximum likelihood and multispecies coalescent trees of single-copy genes showed well-supported relationships for major lineages, which were generally consistent across analyses and genomes, and with previous taxonomic and phylogenetic findings. However, they also revealed high levels of both nuclear and cyto-nuclear discordances. The time-calibrated phylogeny of target-capture data supports an early Miocene origin for Loliinae and Mid-Late Miocene splits for its main broad-leaved and fine-leaved lineages, while current species-rich groups radiated in Plio-Pleistocene times. Hybridization and topological incongruence tests between the nuclear single-copy genes and plastome-based trees using multiple approaches and different sampling subsets confirmed the rampant introgression experienced by Loliinae at deep and shallow nodes. However, hybridization rates differed from lineage to lineage within the major clades and were not correlated with time or ploidy but rather depended on the “hybridogenous” nature of particular lineages. Our analyses detected high hybridization rates in four broad-leaved (Subulatae-Hawaiian, Tropical-South African, Mexico-Central-South American and Leucopoa p.p.) and five fine-leaved Loliinae lineages (American II, Aulaxyper, Afroalpine, American-Neozeylandic) containing rogue species that probably originated from trans-clade crosses and are more likely to hybridize greatly. In contrast, they recovered low hybridization rates in four broad-leaved (Schedonorus-Lolium, Subbulbosae, Drymanthele-Pseudoscariosae-Lojaconoa, Leucopoa pp) and six fine-leaved lineages (Festuca, Psilurus-Vulpia, American I, Exaratae-Loretia, American-Vulpia-Pampas, Eskia), with species derived from single ancestors that hybridize only with close congeners. The levels of intragenomic nuclear discordance could have been magnified by the prevalence of allopolyploids in the Loliinae and by methodological bias in the selection of orthologs; however, our nuclear and plastome trees have revealed key hybrid origins in these grasses. [allopolyploidization, lineage-specific hybridization rates, nuclear and cyto-nuclear discordances, *Festuca* and related genera, phylogenomics, nuclear single-copy genes, plastome].

## Introduction

Reconstructing the phylogenetic relationships of large groups of organisms is challenging due to the existence of biological and genomic events, such as hybridization and polyploidization, gains and losses of genome fractions, and lateral gene transfers, which prevent or confound inferences retrieved from simple bifurcating trees (Marcussen et al., 2015; Dunning et al., 2019; Liu et al. 2019). The complexity is exacerbated in highly reticulate plant lineages where the number of speciation events resulting from introgressions and auto- or allopolyploidizations (and subsequent diploidizations) have occurred frequently and recurrently (Soltis et al. 2016; Mandáková and Lysak 2018; Sancho et al. 2022). Although the construction of evolutionary species networks from multilabelled trees is considered a suitable alternative to incorporate potential hybridizations and allopolyploidizations in the phylogeny (Huber et al. 2006), successfully finding the correct network would depend on whether the available set of multilabelled gene trees contains all the homeologs representing all subgenomes of the allopolyploid (Marcussen et al. 2015). However, incomplete homeologous sampling is common in ancient polyploids in which many copies of duplicate genes have been removed due to genome re-structuring and excess genetic load (Michael 2014), and even in recent polyploids in which chromosomal instability has promoted rapid genome rearrangements and gene losses (Chen and Ni 2006). The use of coalescence-based methods and models applied to multigenomic data that take into account the potential existence of incomplete lineage sorting (ILS), introgression, and gene duplication has been considered a straightforward approach to infer the true phylogeny of organisms at various taxonomic levels (Baker et al. 2022). Gene target capture of hundreds of genes has emerged as a useful tool for building phylogenies for non-model plant groups with limited genomic information and for taxa that are only known or available in historical collections (Johnson et al. 2019; Pérez-Escobar et al. 2021; Thomas et al. 2021; Zuntini et al. 2021; Baker et al. 2022). The recent development of methods that identify and estimate incongruences and conflicts in gene-tree species trees and cyto-nuclear discordances allow researchers to disentangle the intricate evolution of reticulate groups and test hypotheses on their captivating high morphological diversification (Pérez-Escobar et al. 2016; Minh et al. 2020a; Hendriks et al. 2023).

The Loliinae subtribe constitutes one of the most diversified lineages of cool seasonal pooid grasses (Catalán 2006). It consists of more than 600 species of pasture and forage plants, some of which are of considerable ecological and economic importance (e.g., meadow, tall, red and sheep’s fescues, and ryegrasses) or are dominant species in their grassland ecosystems. Loliinae are distributed in temperate and tropical mountainous regions of all continents except Antarctica (Minaya et al., 2017; Moreno-Aguilar et al., 2020; Moreno-Aguilar et al., 2022a). The subtribe comprises the large genus *Festuca,* with more than 500 species, plus 13 closely related genera, totalling about 50 species (Catalán, 2006; Catalán et al., 2007; Moreno-Aguilar et al., 2022b). *Festuca* and four genera (*Megalachne*, *Micropyropsis*, *Podophorus*, *Pseudobromus*) contain perennial species, and the remaining 9 consist exclusively of (*Ctenopsis, Dielsiochloa, Hellerochloa, Micropyrum, Narduroides, Psilurus, Vulpia, Wangenheimia*) or predominantly (*Lolium*) of annual species. Loliinae species show a uniform chromosome base number of *x* = 7 and different ploidy levels, ranging from diploid (2n=2x=14) to tetradecaploid (2n=14x=98) (Martínez-Sagarra et al. 2021).

Hybridization and polyploidy are common in *Festuca*, whose species are mostly allopolyploid (Catalán 2006; Šmarda et al. 2008). *Festuca* species from the southern hemisphere are exclusively polyploid (Dubcovsky and Martínez, 1992; Catalán, 2006; Šmarda et al., 2008; Moreno-Aguilar et al., 2022b). This geographic pattern is consistent with Dispersal Extinction Cladogenesis (DEC) biogeographic models that suggest that the two major lineages of Loliinae, broad-leaved (BL) and fine-leaved (FL), originated in the northern hemisphere, where diploids and polyploids currently coexist, and later colonized the southern continents (Minaya et al. 2017). *Festuca* species have been classified into 12 subgenera, according to Alexeev’s worldwide taxonomic treatment, and some of them into various sections and subsections according to various authors, while some species have not been yet ranked taxonomically (Supplementary Table S1; Catalán et al., 2007; Moreno-Aguilar et al., 2022b, and references therein). Previous phylogenetic studies based on few nuclear (ITS, *b- amylase*, *GBSS*) and plastid (*trn*TL, *trn*LF) loci (Inda et al. 2008; Díaz-Pérez et al. 2014; Minaya et al. 2015, 2017) provided a primary evolutionary framework of Loliinae. These studies demonstrated that *Festuca* is paraphyletic and that Loliinae divides into two main BL and FL lineages, each with several sub-lineages recovered in nuclear and plastid-based topologies. More recent phylogenetic work using genome skimming approaches inferred Loliinae trees from plastome and nuclear 35S and 5S rDNA sequences that were generally consistent with earlier phylogenies, but detected more topological incongruences for some specific subclades, and also expanded the sampling of new lineages of Loliinae (Moreno-Aguilar et al., 2020; Moreno-Aguilar et al., 2022b). Evolutionary investigation of the Loliinae repeatome inferred a consensus network topology that was highly congruent with that of the nuclear 35S rDNA tree, suggesting a plausible scenario of recurrent allopolyploidizations followed by some diploidizations that generated large genome sizes of BL diploids, significantly higher than those of the FL diploids (Moreno-aguilar et al. 2022). Despite these advances, the phylogenomic information of Loliinae based on a substantial sample of single-copy nuclear genes and its potential congruence with data from plastomes, nuclear rDNA, and repetitive elements remain to be evaluated and tested.

We present here the first phylogenomic study of Loliinae based on target capture data generated with the Angiosperms353 nuclear probe set (Johnson et al., 2019). Together with whole-plastome data, we aim to investigate the impact of ancient and recent hybridizations in the evolutionary history of Loliinae and test whether levels of introgression were related to time, ploidy level, or phylogenetic group. Specifically, we aimed to assess whether hybridization occurred similarly in BL and FL Loliinae or preferentially in one of these major clades, and whether some particular intra-clade lineages were more hybridogenic than others.

## Materials and Methods

### Sampling

Samples of 132 species from eight genera of Loliinae (*Festuca*, 118; *Lolium*, 5; *Vulpia*, 4; *Micropyropsis*, 1; *Megalachne*, 1; *Wangenheimia*, 1; *Psilurus*, 1; *Hellerochloa* 1) and three related outgroup grasses (*Oryza sativa, Brachypodium distachyon, Hordeum vulgare*) were included in the study (Supplementary Table S1; Supplementary Fig. S1). The 132 Loliinae samples were first analyzed to capture single-copy nuclear gene targets, 79 of them were analyzed for plastome data for the first time, and 30 were not previously included in phylogenetic studies (Supplementary Table S1). Sampling was carried out from fresh materials collected in the field, collections of silica-dried leaves, and herbarium vouchers (Supplementary Table S1). Plastome data retrieved from 49 previously studied taxa (Moreno-Aguilar et al. 2022a; Moreno-Aguilar et al. 2022b) were incorporated into the analysis (Supplementary Table S1). The selected taxa represent the 28 evolutionary lineages currently recognized within Loliinae (Minaya et al., 2017; Moreno-Aguilar et al., 2022b).

### Genome sequencing

Genomic library preparation and targeted enrichment of Loliinae single-copy nuclear genes were performed with the Angiosperms353 target capture kit at Arbor Biosciences (Michigan, USA), following the protocols outlined in Johnson et al. (2019).

Hybridization capture was performed in pools of 12 libraries following the myBaits v5 manual (https://arborbiosci.com/mybaits-manual/). Capture reactions were pooled in equimolar ratios to form a sequencing pool, which was sequenced on a partial lane of the Illumina NovaSeq 6000 platform in paired-end (PE) mode (2 x 150 bp). Output reads from target enrichment sequencing were checked with FastQC (https://www.bioinformatics.babraham.ac.uk/projects/fastqc/) and trimmed with Trimmomatic (Bolger et al. 2014) preserving reads with a minimum length of 50bp (minlen:50) to reduce the risk of potential misalignments of short reads to genes and trimming bases at the end of the reads with low quality (trailing:30).

Genome skim sequencing was performed on 79 Loliinae DNA samples (Supplementary Table S1). Genomic sequencing of a multiplexed pool of PCR-free KAPA libraries was performed on a HiSeq4000 or HiSeq 2500 (TruSeq SBS Kit v4, Illumina, Inc) in PE mode (2 x 101 bp) as described in Moreno-Aguilar et al. (2020). Illumina PE reads were quality-checked using FastQC and adapters and low-quality sequences were trimmed and removed with Trimmomatic. Loliinae genomic samples used in subsequent analysis contained between 1.5 – 35.0 million reads (average 16.0 million reads) with insert sizes ranging between 69 – 260 bp (Supplementary Table S1).

### Sequence assemblies of target capture data and hybrid-polyploid detection (AD and LH metrics)

Loliinae single-copy nuclear genes were recovered from the target-enriched PE reads using the HybPiper v.1.3.1 pipeline (Johnson et al. 2016). This was done by mapping the filtered reads against the template sequences of the 353 low-copy nuclear genes (available at https://github.com/mossmatters/Angiosperms353) using the Burrows-Wheeler Alignment (BWA v.0.7) (Li et al. 2009), and then through *de novo* assembly of mapped reads for each gene separately using SPAdes v. 3.13 (Bankevich et al. 2012), with the default minimum coverage threshold of 8× (Supplementary Fig. S2). The HybPiper script *retrieve_sequences.py*, which generates a single sequence per gene that is selected based on length, similarity, and depth of coverage criteria, was used to obtain the strict data set from Loliinae, made mostly of nuclear single-copy genes (scg-strict data set) (Appendix 1a). HybPiper selects the allele with the best coverage and recovery for a gene based on the relative nucleotide frequency of each heterozygous site (Hendriks et al. 2022). To identify and extract putative paralogs from this data set, we consecutively run the *paralog_investigator.py* and *paralog_retriever.py* scripts in HybPiper to obtain the paralog fasta files. These paralogs were discarded from the Loliinae scg-strict data set through phylogenetic filtering, thus keeping only the optimal ‘single-copy’ allele per gene and per sample for final phylogenetic reconstruction (Supplementary Table S2; Appendices 1a, 1b).

For the HybPiper data set, individual genes were aligned with MAFFT v.7.490 (Katoh & al., 2002) using the iterative refinement method *--maxiterate* 1000. Empty genes for any Loliinae sample were removed from downstream analysis. Phyutility 2.2.6 (Smith & Dunn, 2008) was used to remove sequences with insufficient coverage (<30%, *-clean* 0.3) in well-occupied columns of each gene alignment. Gene alignments were visually inspected with Geneious Prime to detect potentially misaligned sequences. Sequences with less than 60% of the total alignment length were removed to enhance the quality of the data sets, filtering out species with potentially low phylogenetic information. This process generated the respective multiple sequence alignments (MSAs). To check alignment quality, we estimated summary statistics of gene alignments using AMAS v.0.98 (Borowiec, 2016). These statistics included alignment length, missing data, and number of parsimony informative sites (Appendix 1a).

We used HybPhaser v2.0 (Nauheimer et al. 2021) to detect potential hybrids in our target capture data set of presumably highly hybridogenic Loliinae. This tool maps raw sequence data to the HybPiper contigs taking into account SNP variation using nucleotide ambiguities and quantifying divergence between gene variants. The gene variants correspond either to paralogs or to hybrids (“homeologs” in allopolyploids; Sancho et al. 2022) (Supplementary Table S3). Samples showing high SNP content in genes are likely hybrids or polyploids and are phased for the different copy variants (alleles) into phased haplotypes after discarding putative paralogs (>1.5× the interquartile range above the third quartile of mean SNPs) (Nauheimer et al. 2021; Hendriks et al. 2022). We ran HybPhaser to detect potential hybrid/polyploid samples and generated a data set of phased accessions of Loliinae (scg-inclusive data set) (Supplementary Table S3). We used R scripts to compute two metrics indicative of hybridization, sample allele divergence (AD, percentage of SNPs in all genes) and locus heterozygosity (LH, percentage of genes with SNPs). These values reflect the hybridization history of the samples (Nauheimer et al. 2021; Hendriks et al. 2023). We classified the Loliinae samples into three LH-vs-AD hybridogenic classes (diploid: low-to-high LH and low AD; old polyploid: medium LH and high AD; high polyploid: high LH and high AD) (Hendriks et al. 2023).

### Sequence assemblies of plastomes

Whole-plastome sequences for 79 new Loliinae samples were assembled from their respective genome skimming PE reads, following the procedures indicated in Moreno-Aguilar et al. (2022b). Plastome assembly was performed with Novoplasty v.2.7.1 (Dierckxsens & al., 2017) using the plastome of *Festuca pratensis* (JX871941) as reference sequence and standardized parameters (k-mer: 30-39, insert size: ∼69-200 bp, genome range: 120,000–140,000 bp, and PE reads: 101-150 bp). Furthermore, to recover plastome sequences in data with a low number and quality of total PE reads, plastome assembly was performed using a read-mapping strategy to, respectively, closely related *Festuca* plastomes using Geneious Prime (Supplementary Table S1). The data for 49 representative Loliinae taxa from previous studies (Moreno-Aguilar et al., 2020; Moreno-Aguilar et al., 2022a, 2022b) were incorporated into, existing plastome data set (Supplementary table S1). Whole plastomes sequences were aligned separately with MAFFT v.7.031b, and trimAl v. 1.2rev59 (Capella-Gutiérrez et al. 2009) was used to remove poor quality regions from the MSA by enforcing the -*automated*1 parameter.

### Phylogenomic reconstructions and intragenomic nuclear discordance

We used three different approaches to analyze our genomic data sets and reconstruct the Loliinae phylogenies. First, trimmed MSAs of the Loliinae scg-strict data set were used for a maximum likelihood (ML) supermatrix approach using IQtree2 v. 2.2.2.6 (Nguyen & al., 2015, Minh et al. 2020a), concatenating all loci and using the ‘edge proportional’ partition model with single loci as partitions and model selection implemented via ModelFinder (Kalyaanamoorthy & al., 2017) for each partition. Branch support was calculated from 1000 UltraFast Bootstrap replicates (Hoang & al., 2018) and gene (gCF) and site concordance factors (sCF, parameter –scf 1,000; Minh et al. 2020b) for all nodes. Second, individual ML trees of each nuclear single-copy gene, as well as ML trees of the whole plastome MSA were also computed separately using the same IQtree2 procedure. *Oryza sativa* and *Brachypodium distachyon* (plus *Hordeum vulgare*, single-copy gene dataset) were used to root all the trees. Third, to account for potential ILS events between closely related Loliinae lineages and putative topological incongruences between single-copy nuclear gene trees, we inferred a species tree by analyzing the Loliinae scg-strict data set under multispecies coalescent (MSC) using ASTRAL-III v.5.7.8 (Zhang et al., 2018), which was fed with the individual IQtree2 ML gene trees. All gene trees were rooted using *Oryza sativa, Brachypodium distachyon,* and *Hordeum vulgare,* with the *pxrr* function in Phyx (Brown & al., 2018), except four trees that were rooted using the early diverging lineages of *Festuca lasto* and *F. drymeja* as outgroup. Branches with likelihood bootstrap support values <30% in gene trees were collapsed using *nw_ed* from Newick Utilities 1.6.0 (Junier and Zdobnov, 2010). To estimate the intragenomic discordance in the nuclear dataset, we analyzed in R the normalized quartets scores generated by ASTRAL-III for the main topology and the first and second alternative topologies when inferring the MSC species tree. Although the MSC model is only consistent when ILS is the only source of gene tree discordance, comparison with the supermatrix ML tree can identify highly compatible clades (Nauheimer et al. 2021).

### Evaluation of cytonuclear topological incongruence, phylogenetic signal, hybridization tests, and dating analysis

To assess the degree to which the phylogeny retrieved by the nuclear coding genome tracked that of the organelle (plastome) genome, we also applied the Procrustean Approach to Cophylogenetics (PACo) procedure implemented in R (Balbuena et al. 2013; Pérez-Escobar et al. 2016) to the nuclear scg-strict supermatrix ML trees versus the plastome supermatrix ML trees (bootstrap trees) of 128 common Loliinae species. Due to the high degree of intragenomic congruence of the Loliinae plastome-coding genes (Minaya et al. 2017; Moreno-Aguilar et al. 2020), whole-plastome alignment was employed to generate 1,000 bootstrap plastome trees. To do this, a phylogenetic reconstruction was carried out using IQtree2, providing the program with partition information for each nuclear and plastome alignment, executing 1,000 ultrafast bootstrap replicates and saving the respective bootstrap trees (ufboot). This procedure evaluates the similarities between any pair of topologies by comparing the Euclidean distances that separate the terminals in both trees through the Procrustean superposition (Balbuena et al., 2013). The sum of the squared residuals (the disparity between an observed value and a fitted value derived from a model) for each association and each pair of topologies evaluated can be interpreted as a concordance score because it is directly proportional to the magnitude of the topological conflict for the pair of terminals considered (Pérez-Escobar et al. 2016). Differences in terminal position between nuclear and plastome ML trees were summarized in bar plots using the R package *ggplot2* (Wickham 2016). The sum of squared residuals for each pair of nuclear and plastome gene terminals was classified into quartiles, and the magnitude of discordance was assessed by the proportion of genes binned in quartiles 3 and 4 (50% and 75%) in each terminal. The higher the number of genes binned in quartiles 3 and 4, the greater the discordance is (Pérez-Escobar et al. 2021). Additionally, we contrasted the congruence of nuclear vs nuclear and nuclear vs plastome topologies using the approximately unbiased (AU) test (Shimodaira 2002) implemented in IQTree2.

To search for overall evidence of hybridization within Loliinae, we applied HyDe v0.4.3 (Blischak et al. 2018) which detects hybridization using phylogenetic invariants under the coalescent model with hybridization, and tests sets of triples across the whole phylogeny. The supermatrix of 234 genes (scg-strict data set) was concatenated using FASconCAT-C master software and used as input for HyDe. A high frequency of gamma scores (hybridization proportion) indicates a high prevalence of hybridization in all ingroup taxa.

We dated the origins of the Loliinae lineages using the concatenated single-copy gene IQtree ML phylogeny with branch length information and the treePL software (Smith and O’Meara 2012). We constrained four nodes of the tree using dates inferred from our previous studies. Three nodes were calibrated using secondary age constrains for the crown nodes of the BOP (Oryza + Pooideae) clade (normal prior maximum = 55.08, minimum = 47.76 Ma), Loliinae clade (normal prior maximum = 23.13, minimum = 16.02 Ma), BL Loliinae clade (normal prior maximum = 20.45, minimum = 12.15 Ma), and a fourth node was calibrated using a minimum age constrain for the crown node of fine-leaved Loliinae (lognormal prior maximum = 19.75, minimum = 14.13 Ma) based on a *Festuca* sect. *Festuca* leaf macrofossil dated to the Early Miocene (Moreno-Aguilar et al., 2020). The analyses were performed using smoothing values of 0.1 to estimate the best optimization parameter values (3-3-5), which were then used in a subsequent analysis to calculate divergence times.

## Results

### Sequencing data

On average, 2.95 M PE reads per sample, ranging from 1.29 M in *F. pyrenaica* to 8.64 M in *F. californica*, were generated for target capture data. The average recovery of reads per genus was similar for *Festuca* (3.05 M), *Vulpia* (3.35 M), *Wangenheimia* (3.31 M), *Micropyropsis* (3.69 M), and *Psilurus* (3.62 M), and slightly less for *Megalachne* (2.59 M) and *Lolium* (2.42 M) (Supplementary Tables S1, S2; Appendices 1a, 1b). The total number of Loliinae nuclear target genes obtained from the Angiosperms353 kit was 351. The mean number of genes recovered by HybPiper with >50% of the target length for Loliinae samples across the entire sampling was 252, representing 71.30% of the target loci, while the mean number of genes with >75% of the target length was 175 (49.56%) (Supplementary Table S2; Appendices 1a, 1b). We detected 107 potential paralog instances via HybPiper’s paralog warnings. After excluding paralogous copies in the Loliinae scg-strict data set and three genes with sequences for fewer than 30 species, the final data set consisted of 234 genes and 132 species, representing on average 68.27% of the 353 reference genes. Individual single-copy nuclear gene alignments ranged from 75bp to 3,335bp, with a mean length of 659bp (Supplementary Table S2, Appendices 1a, 1b). The final alignment of 234 concatenated and partitioned nuclear genes for 132 species in the scg-strict data set was 158,926 bp in length. The percentage of missing data was 25.88% and the percentage of parsimony informative sites 29.5% (45,953bp) (Appendix 2). The number of loci retrieved by HybPhaser for the Loliinae ranged from 209 (*F. lugens*) to 327 (*F. parvigluma*), representing, respectively, 27.8 to 68.2 of the total (345 genes) (Supplementary Table S3).

Genome skimming data from 79 newly sequenced samples ranged from 1,56 M (*F. livida*) to 35,89 M (*F. glauca*) PE reads. The newly assembled complete plastomes ranged between 103,079 (*F. livida*) and 134,746bp (*F. costata*), which is consistent with plastome length values obtained in previous studies of Loliinae for the respective FL and BL clades (Moreno-Aguilar et al., 2022b) (Supplementary Table S1). Most of the newly assembled plastomes showed good read coverage (>40×). The MSA of the complete plastomes was 135,487 bp in length [19,594 (14.5%) variable sites, 8164 (6.02%) parsimony informative sites], with *F. livida, V. muralis* and *P. incurvus* being the samples with the highest percentage of missing data (23%) (Appendix 2). The newly obtained sequences from each data set were deposited in GenBank (Supplementary Table S1).

### Loliinae nuclear target capture phylogenies, intragenomic discordances, and detection of hybrids and polyploids

Phylogenetic trees recovered from the Loliinae scg-strict data set using supermatrix ML and MSC approaches showed topologies that were relatively highly congruent with each other (Figures 1, 2a; Supplementary Fig. S3). Both phylogenies supported the major split of BL and FL Loliinae clades, as well as divergences from most of the BL (Drymanthele-Phaeochloa + Scariosae + Lojaconoa + Pseudoscariosa; Tropical–South Africa; Subbulbosae + Leucopoa, Schedonorus) and FL lineages (Eskia; American II; American-Neozeylandic; American I; Psilurus-Vulpia(px); Festuca + Wangenheimia; Afroalpine). However, the two trees showed conflict regarding the inferred relationships of some BL (Mexico-Central American-South American (MCSA) I and II) and FL (‘intermediate’ Subulatae-Hawaiian; American–Vulpia Pampas; Loretia + Exaratae; Aulaxyper) lineages (Supplementary Fig. S3). The main topological discordances were related to the different locations of the BL MCSA I (Glabricarpae, Asperifolia, Drymanthele s. l.) and MCSA II (Erosiflorae, Ruprechtia, Coironhuecu) groups, forming an intermediately evolved sister clade of Subbulbosae-Leucopoa / Schedonorus in the ML tree (Figure 1), but resolved as sister to an ancestral Drymanthele-Phaeochloa + Scariosae + Lojaconoa + Pseudoscariosa / Tropical-South Africa clade in the MSC tree (Figure 2a; Supplementary Fig. S3). In both trees, all MSCA lineages descended from a common ancestor except *F. argentina* (Coironhuecu), which nested within the Subbulbosae + Leucopoa clade and was resolved as sister of *F. altaica* (Figures 1, 2a). The Exaratae-Loretia group was divided into a clade ((*V. membranacea* / *V. sicula*), *F. plicata*) sister to Psilurus-Vulpia (px), and other lineages (*F. hephaestophila, F. capillifolia* / *F. pyrenaica*) which showed different relationships with Aulaxyper p. p. lineages and to Afroalpine or to American-Vulpia-Pampas and American I in the ML and MSC trees (Figures 1, 2a; Supplementary Figure S3). Our largest sampling of taxa (Supplementary Table S1) enriched the phylogenetic circumscriptions of several clades or groups with newly studied species (Festuca: *F. gracilior, F. kolesnikovii, F. marginata, F. mollisima, F. reverchonii, F. yvesii*; Aulaxyper: *F. raddei, F. richardsonii*; Australia-Tasmania: *F. plebeia, F. asperula*; American-Vulpia-Pampas: *F. samensis*; American-Neozeylandic: *F. kurtziana*; American I: *F. imbaburensis*; American II: *F. carazana, F. dasyantha, F. distichovaginata, F. dolichophylla, F. glumosa, F. humilior, F. laegaardii, F. monguensis, F. procera, F. parciflora, F. rigidifolia, F. setifolia, F. sodiroana, F. subulifolia, F. versuta, F. weberbaueri*; Subulatae-Hawaiian: *F. leptopogon*; Eskia: *F. acuminata, F. woronovii*; Asperifolia: *F. lugens*). Most of these species clustered in the same groups in the ML and MCS trees, although their relationships differed in some cases from tree to tree (Figures 1, 2a, Supplementary Figure S3). The strongly supported Australian *F. asperula* / *F. plebeia* clade (new 29^th^ Loliinae lineage) was closely related to the Aulaxyper group, but constituted a separate basal Australia-Tasmania lineage of this grade in both trees (Figures 1, 2a, Supplementary Fig. S3). The taxonomically and phenotypically diverse American II lineage included species classified within the fine-leaved *Hellerochloa* (*H. livida*) and *Festuca* sect. *Festuca* (e. g., *F. andicola, F. orthophylla*) or within different fine-leaved supraspecific *Festuca* ranks (e. g., *F. versuta* (*F*. subgen. *Drymanthele* sect. *Texanae*) / *F. subverticillata* (*F*. subgen. *Obtusae*), *F. flacca* (*F*. subgen. *Subulatae* sect. *Subulatae*) (Supplementary Table S1), although they were all nested in the same clade in both trees. Our ML and MSC analyses indicate that *F. subverticillata* belongs to the American II clade while *F. hephaestophila* is not affiliate with the Festuca lineage but is placed within the Exaratae grade (Figures 1, 2a, Supplementary Fig. S3). The broad-leaved species *F. californica* (*F*. subgen. *Leucopoa* sect. *Breviaristatae*) and *F. subuliflora* (*F*. subgen. *Subuliflora*) were resolved as sister taxa (Figure 1) or closely related species (Figure 2a) in the ML and MCS phylogenies, respectively, nesting in early divergent lineages within the FL clade. *F. muelleri* (*F*. subgen. *Drymanthele* sect. *Banksia*) was strongly resolved as a sister lineage to the Schedonorus clade in both trees (Figures 1, 2a). The Eurasian broad-leaved *F. modesta* (*F*. subgen. *Drymanthele* sect. *Muticae*) and *F. calabrica* and *F. olgae* (*F*. subgen *Leucopoa*), and the South African *F. scabra* fell within an expanded Tropical-South African+ (plus) clade in the ML and MSC trees (Figures 1, 2a, Supplementary Figure S3).

**Figure 1.**
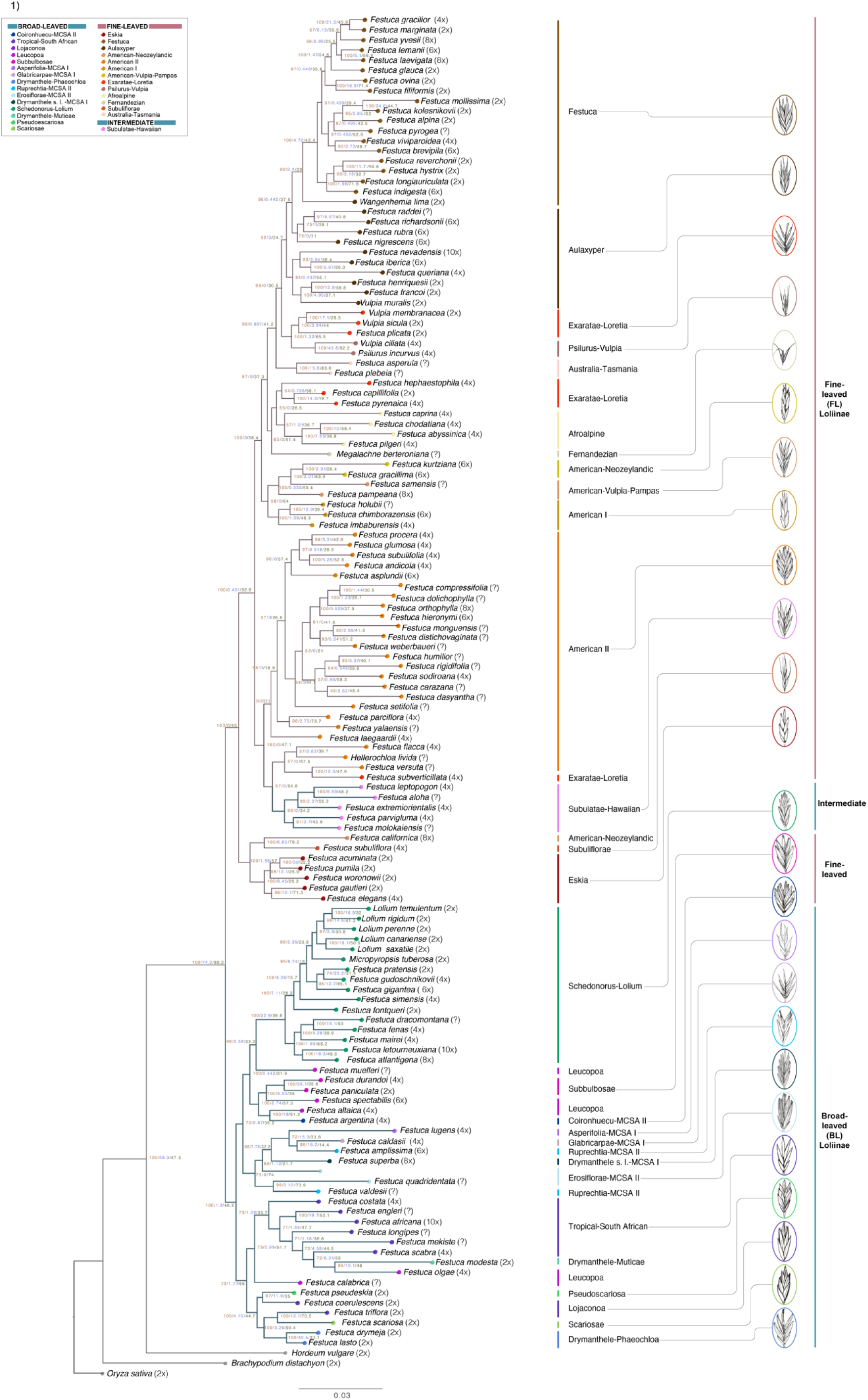
Maximum Likelihood (ML) single-copy gene phylogeny of Loliinae constructed from a concatenated supermatrix (scg-strict data set) of 234 genes and 132 taxa. Numbers on branches indicate UltraFast Bootstrap support (BS, maroon), gene concordance factor (gCF, blue) and site concordance factor (sCF, green) values (see Supplementary Table S4), respectively. Scale bar: number of mutations per site. Color codes of Loliinae lineages are indicated in the chart. Drawings of spikelets are shown for representative species of each Loliinae lineage. Drawings by M. F. Moreno-Aguilar.

**Figure 2.**
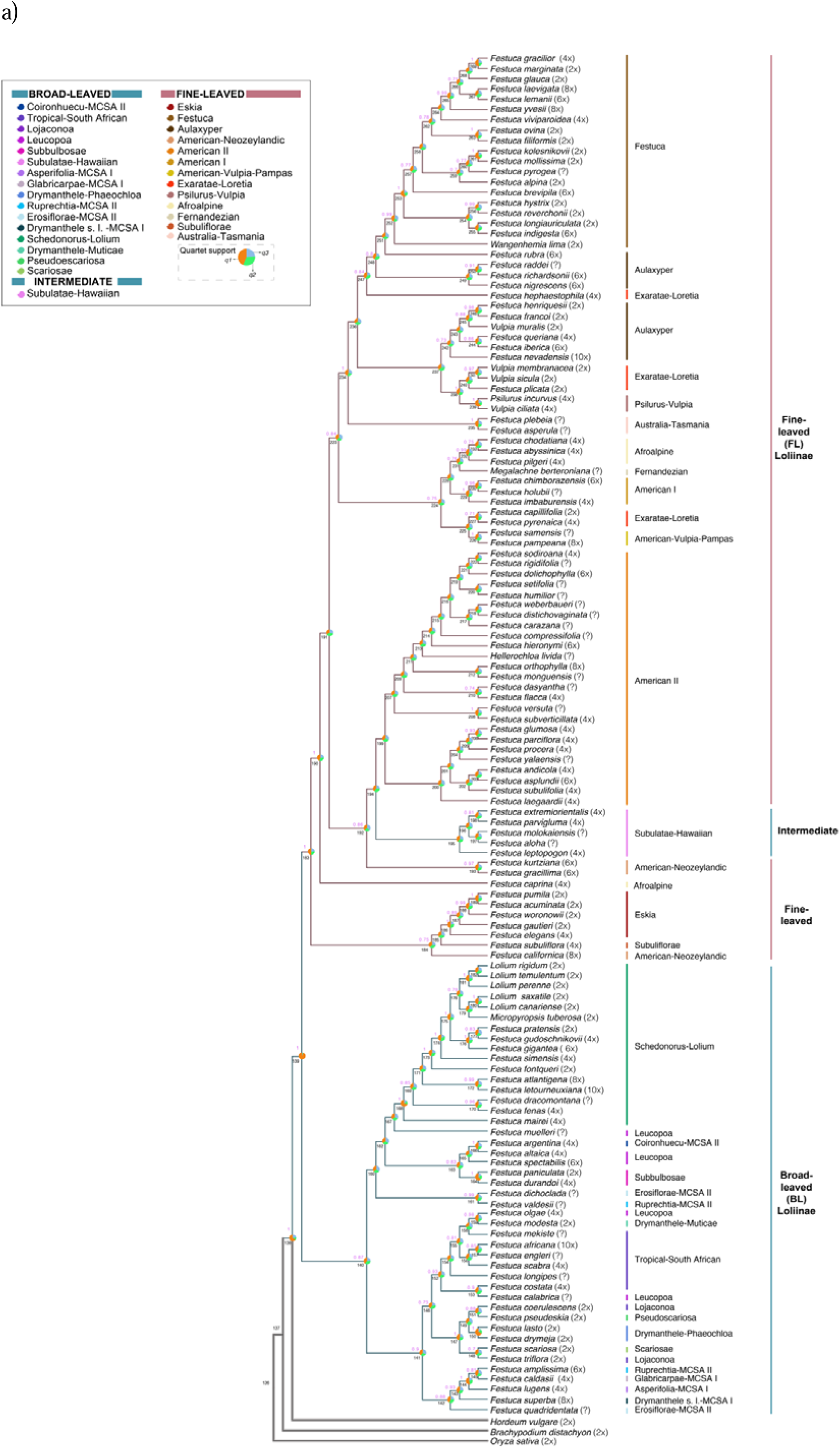
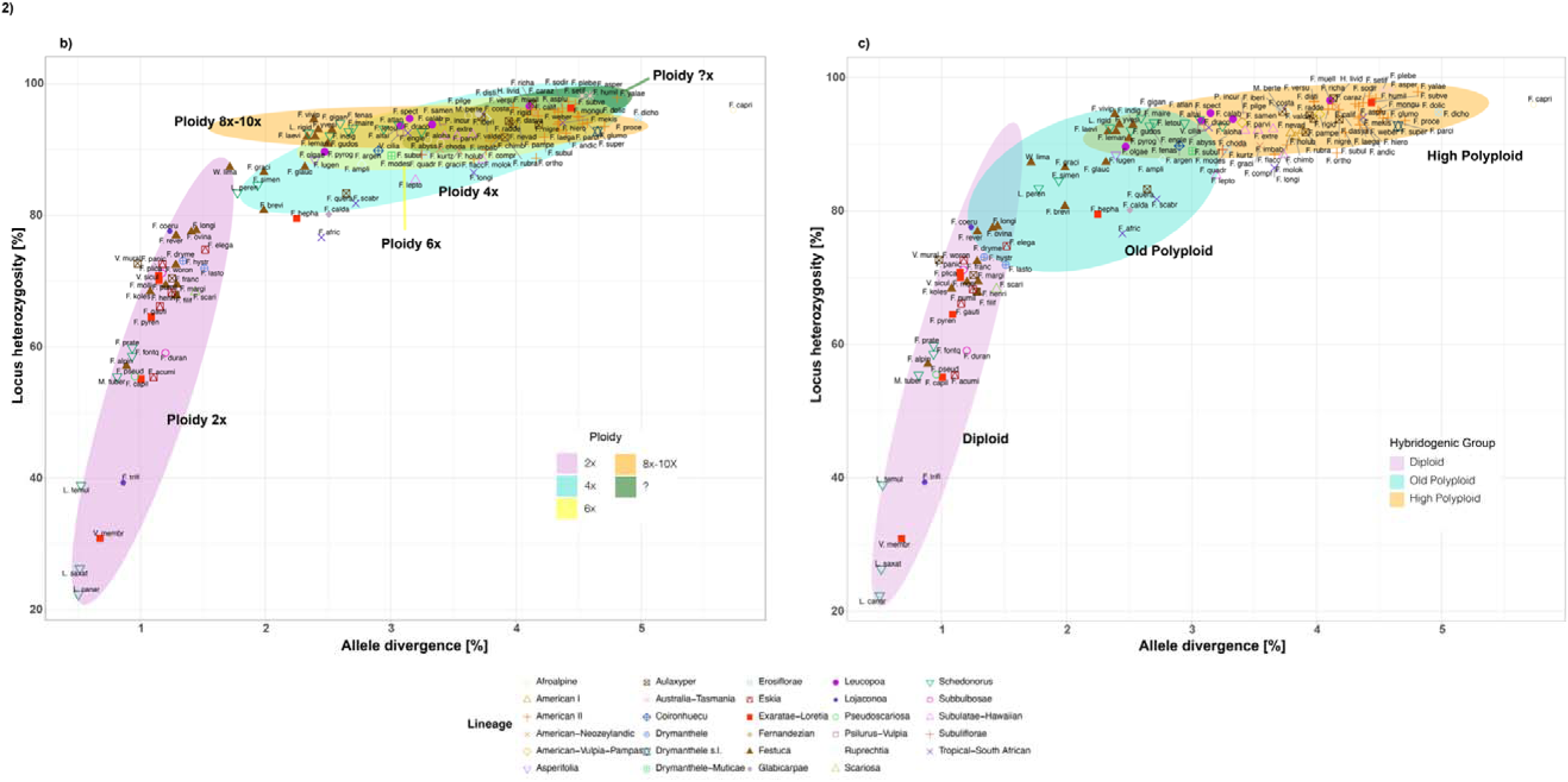
**(a)** Multispecies Coalescent (MSC) ASTRAL species tree of 132 Loliinae taxa inferred from 234 ML single-copy gene trees (scg-strict data set). Numbers above branches show Posterior Probability Support (PPS) values and below their consecutive numbering. Pie diagrams at nodes correspond to quartet support values for branches of the best tree and the alternative topologies (*q1*, *q2*, and *q3*, see color codes in the chart and Supplementary Table S5). *Oryza sativa* was used to root the trees. Color codes of Loliinae lineages correspond to those of Figure 1. **(b, c)** Scatterplots displaying the locus heterozygosity (LH) and allele divergence (AD) values of the studied samples retrieved from HybPhaser. Colored ellipses indicate four main groups of ploidy-levels (b) and three hybridogenic classes (diploid: low-to-high LH and low AD; old polyploid: medium LH and high AD; high polyploid: high LH and high AD) (c). Color codes for ploidy levels and hybridogenic classes, and symbols for Loliinae lineages are indicated in the respective charts.

Node support was high for most tree branches in both the ML (ultrafast median BS values mostly 100%) and MSC (most median PPS values close to 1) trees (Figures 1, 2a). However, the mean percentages of gene concordance factor (gCF) and site concordance factor (sCF) were relatively medium-low, at 41.69% and 3.05%, respectively, in the ML tree (Figure 1, Supplementary Table S4). Similarly, genomic discordance between single-copy nuclear genes used to reconstruct the scg-strict MSC phylogeny of Loliinae was high for most lineages (Figure 2a; Supplementary Table S5). The estimation of the proportion of gene tree quartets concordant with the ASTRAL-III species tree via normalized quartet scores indicated considerable intragenomic incongruence. The branches of the MSC tree obtained values ranging from 30.16 – 94.26% for the main topology (*q1*); 2.79 – 37.54 % for the first alternative topology (*q2*), and 2.95 – 37.22 % for the second alternative topology (*q3*), but with a mean of only 41.46% across all nodes of the species tree (*q1*) (Supplementary Table S5). The highest intragenomic concordances were found for the crown nodes of Loliinae (94.26% *q1*), Schedonorus (71.70%), Drymanthele – Phaeochloa (69%) and Psilurus-Vulpia-px (63%) followed by the moderate concordance of those of Subbulbosae (62.03%), broad-leaved MCSA I-II (42.57%,), FL Loliinae *sensu lato* (including Subulatae-Hawaiian) (38.36%,) and fine-leaved Eskia (38.71%), while the remaining lineages showed very low concordances (Figure 2a, Supplementary Table S5). The AU test found no significant differences between both topologies where the ML tree is considered the best tree (0.824) compared to the MSC tree (0.176). This suggests that other potential causes of discordance, such as introgression, may be causing the incongruence between the gene trees and species trees.

A first estimation of the levels of introgression existing within Loliinae was obtained from our HyDe analysis, which provided gamma scores of 0.3 to 0.7 with high frequency (>400) (Supplementary Fig. S4), indicating the high occurrence of hybridizations throughout the subtribe. To obtain more precise data on hybridization rates in specific Loliinae lineages, we analyzed the HybPhaser data of the scg-inclusive data set using the 345 saved alleles. Estimations of the proportion of locus heterozygosity (LH) and of allele divergence (AD) over all available genes per sample indicated that, although some samples (34) showed low range values, corresponding to non-hybrids (diploids) (LH<60%; AD<0.1), others (98) had old- or high-polyploid signatures (Figures 2b, 2c; Supplementary Table S3, Supplementary Fig. S5). Within the polyploids, we were able to separate 82 samples as high polyploids and 16 as old polyploids (Figures 2b, 2c; Supplementary Table S3, Supplementary Fig. S5). This hybridogenic classification fitted most of the inferred ploidy levels of the studied taxa but with some exceptions (Supplementary Table S3). Afroalpine (100%), American-Neozeylandic (100%), American-Vulpia-Pampas (100%), American I (100%), American II (100%), Aulaxyper p.p. (60%), Australia-Tasmania (100%), Drymanthele s.l. (100%), Drymanthele-Muticae (100%), Erosiflorae (100%), Fernandezian (100%), Leucopoa (80%), Psilurus-Vulpia (100%), Subulatae-Hawaiian (100%), Subuliflorae (100%) and Tropical-South African p.p. (66.6%) were the Loliinae lineages richest in high polyploids, while Asperifolia (100%) and Glabricarpae (100%) had more old polyploids. In contrast, Drymanthele-Phaeochloa (50%), Eskia (20%), Exaratae-Loretia p.p. (7.14%), Lojaconoa (0%), Pseudoscariosa (0%), Scariosae (0%), and Subbulbosae (0%) were the less hybridogenic Loliinae lineages. Festuca and Schedonorus-Lolium presented a relatively balanced representation of diploids and high-polyploids (Festuca: 47.4% and 31.6%; Schedonorus-Lolium: 37.5% and 50%), with Festuca showing also several old polyploids (21%) (Figures 2b, 2c; Supplementary Table S3, Supplementary Fig. S5).

### Plastome phylogeny of Loliineae

The topology of the strongly supported plastome ML tree (Figure 3b, Supplementary Fig. S6) showed general concordance for the major BL and FL clades with that of the single-copy nuclear gene ML tree (Figures 1, 3a), although the composition and the relationships between some lineages differed. In the plastome BL clade, most of the MCSA taxa plus South African *F. scabra* and *F. longipes* and Eurasian *F. calabrica* formed a sister clade to the remaining broad-leaved taxa (Figure 3b). Within this last clade, successive splits separated the lineages Lojaconoa-Pseudoscariosa, Drymanthele (Phaeochloa)-Scariosae, Tropical-South African (plus *F. olgae*), Subbulbosae-Leucopoa (plus *F. valdesii*) and Schedonorus-Lolium. In the plastome FL (*sensu lato*) clade, the “intermediate” American-Neozeylandic lineage (*F. californica* / *F. gracillima*) split first, followed by those of Subuliflorae + Breviaristatae (*F. altaica*) + Eskia p. p. I (*F. acuminata* / *F. pumila*), and Eskia p.p. II (*F. elegans*, *F. gautieri, F. woronowii*) + American I + American II p.p. (*F. monguensis, F. parciflora, F. kurtziana*). The divergence of two sister clades followed, one including the successive splits of the American-Vulpia-Pampas/Fernandezian, Psilurus-Vulpia(px), the Exaratae-Loretia grade, and Subulatae-Hawaiian lineages, and the other the Festuca-Wangenheimia, Aulaxyper-Vulpia(2x), and American II + Afroalpine lineages (Figure 3b, Supplementary Fig. S6).

**Figure 3.**
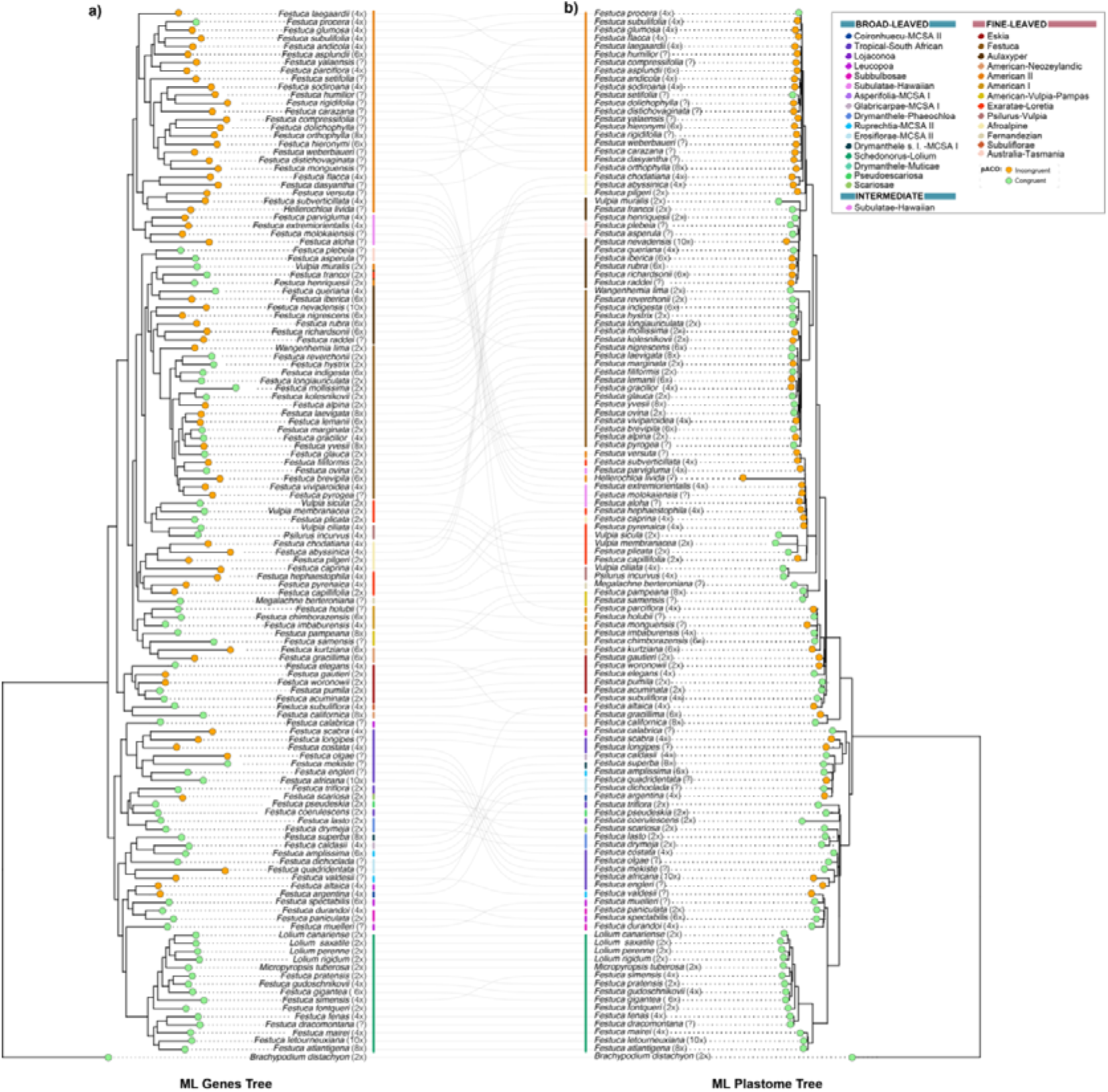
Topological incongruence analysis of the Loliinae single-copy gene ML tree (scg-strict data set) (a) vs whole-plastome ML tree (b) for 128 common samples. The Procrustean Approach to Cophylogeny (PACo) was used along with phylograms obtained from 1,000 bootstrap replicates for each dataset. Orange dots highlight terminals with topological incongruence strongly supported in both nuclear and plastid trees, while green dots indicate congruent terminals. Dashed lines show associations between incongruent terminals. Color codes of Loliinae lineages are indicated in the chart.

### Divergence time estimation analysis

Our penalized likelihood divergence time analysis performed on the single-copy nuclear gene ML tree (Figure 4) provided age estimates for the stem and crown Loliinae nodes dating to the Late-Oligocene (25.76 Ma) and Early Miocene (17.58 Ma), respectively. Early and Mid-Miocene divergences were inferred for the BL (16.3 Ma) and FL (16.06 Ma) ancestors, and Pliocene-to-Quaternary ages for those of the more recently evolved broad-leaved [Drymanthele-Phaeochloa + Lojaconoa + Scariosae + Pseudoscariosa (14.73 Ma), Tropical – South Africa + Drymanthele-Muticae + Leucopoa (*F. olgae, F. calabrica*) (13.83 Ma), MCSA (14.1 Ma), Subbulbosae + Leucopoa (11.67 Ma), Schedonorus (9.62 Ma)] and fine-leaved [Eskia + Subuliflorae Breviaristatae + American-Neozeylandic (*F. californica*) (13.81 Ma), Subulatae – Hawaiian (10.52 Ma), American II (12.37 Ma); American I + American-Neozeylandic + American-Vulpias-Pampas (12.58 Ma), Fernandezian (12.49 Ma), Afroalpine (11.42 Ma), Exaratae – Loretia p. p. (11.44 Ma), Aulaxyper – Vulpia (10.68 Ma), Festuca – Wangenheimia (8.76 Ma)] lineages (Figure 4).

**Figure 4.**
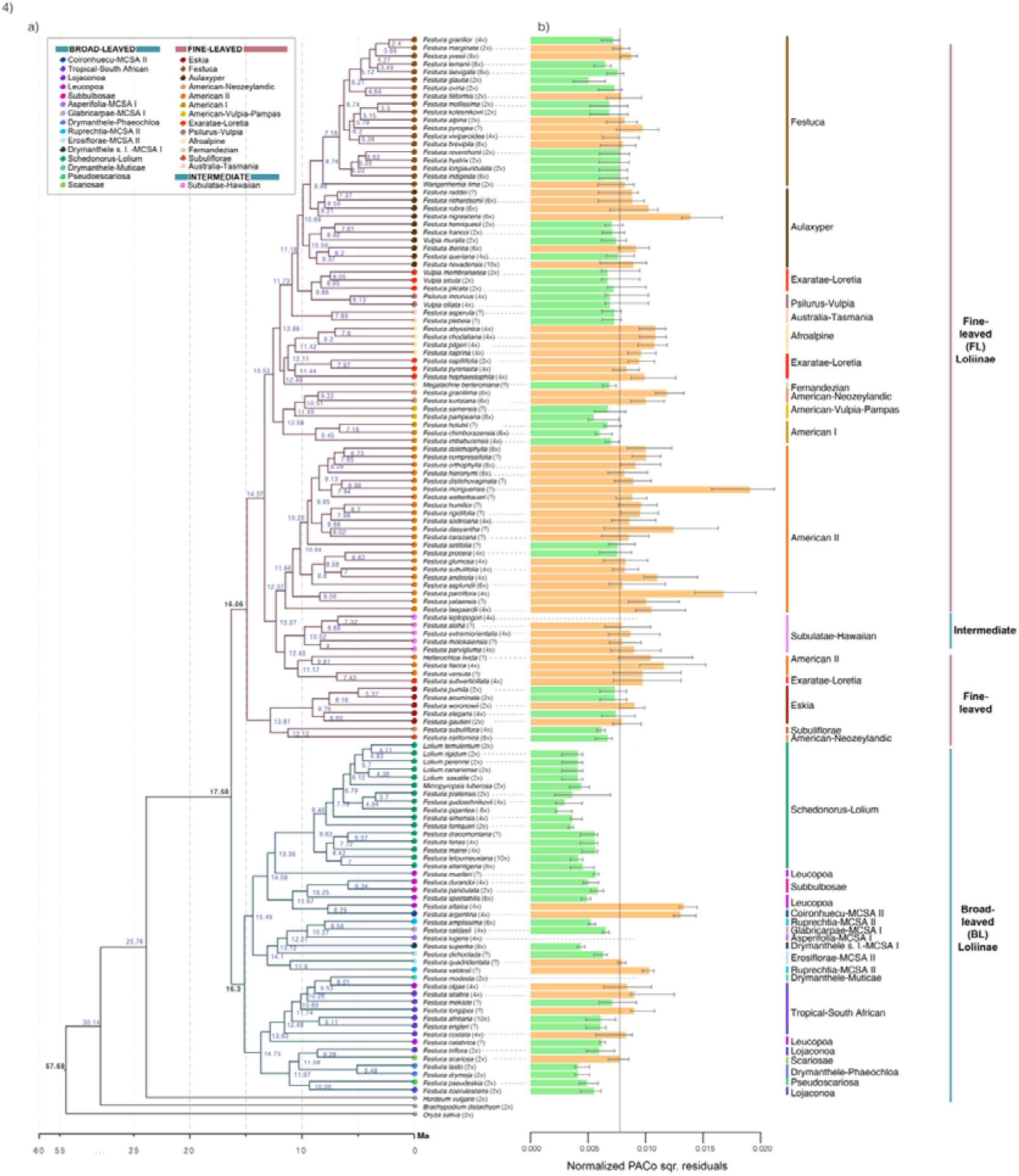
**(a)** Maximum penalized likelihood dated tree constructed with TreePL using 234 nuclear single copy genes of 132 Loliinae taxa and four calibrations. Nodal values show the inferred divergence ages; stars indicate the calibrated nodes. **(b)** Boxplot showing PACo normalized squared residual values obtained from 1,000 random replicates of nuclear-plastome associations. The horizontal black line equals 1/n=0.0077, where *n*= 128 is the number of common nuclear-plastome terminal-associations; median values above this threshold are expected to be linked to species that show incongruence between nuclear and plastome-derived trees (orange boxes) while values below the threshold correspond to topologically congruent species (green boxes) (see also Supplementary Fig. S5).

### Cytonuclear topological incongruence tests and levels of hybridization

AU pairwise topological congruence testing between the nuclear supermatrix ML tree and the plastome ML tree showed that each topology did not differ significantly (p<0.001) from the other. However, topological incongruence analysis performed using the procrustean approach to cophylogeny on the single-copy nuclear gene ML trees vs plastome ML gene trees suggested 61 terminals as potentially conflicting (Figures 3, 4b, Supplementary Fig. S5). The squared residual values of these terminals, calculated individually for each nuclear gene tree and with 46% of their values assigned to quartiles 3 and 4, were generally higher compared to 67 non-conflicting terminals (Figure 4b, Supplementary Fig. S5).

## Discussion

### Limitations of phylogenetic analyses in Loliinae: advantages and disadvantages of single-copy nuclear genes

Our evolutionary analysis has provided the first phylogeny of Loliinae based on hundreds of nuclear-coding genes (Figures 1, 2a, 3a; Supplementary Fig. S3). The retrieved phylogeny of the 234 single-copy genes (scg-strict data set) shows high resolution and strong support for the BL and FL Loliinae lineages in the concatenated ML tree (Figure 1). This topology is in general agreement with the topology inferred from the plastome (Figure 3b, Supplementary Figure S6) for the major lineages of broad and fine-leaved Loliinae, as well as with earlier phylogenies based on a few nuclear and plastid loci (Inda et al. 2008; Díaz-Pérez et al. 2014; Minaya et al. 2015, 2017) and nuclear repetitive elements (Moreno-Aguilar et al., 2022a). We can, therefore, conclude that compartments of the nuclear (nDNA) and organellar (pDNA) genomes reconstruct a congruent evolutionary scenario for the divergences of the main lineages of Loliinae, which allows the phylogenetic and statistical test of specific hypotheses about their potential origins.

However, branch support decreases and some relationships differ between recently evolved BL and FL Loliinae sub-lineages in the single-copy gene MSC ASTRAL-III species trees (Figure 2a). The topological incongruences detected between the concatenated partitioned ML tree and the ASTRAL species tree can be attributed to the severe impact of ILS on the evolutionary history of the youngest Loliinae groups (e.g., BL MCSA and FL American II taxa; Figures 1, 2a, 4a, Supplementary Figure S3). In particular, the value of the quartets scores for the main topology of the ASTRAL-III species tree was generally moderate to high (30.16 – 94.26%) relative to its first and second alternative topologies (Figure 2a; Supplementary Table S5). For several of the ingroup nodes (41.46%), the *q1* topology had the largest quartets scores, indicating that a majority of the possible gene trees were concordant between them, while for several nodes the alternative topologies (*q1, q2, q3*) had similar values (∼0.37), which reflects that the possible gene trees were present with almost the same frequency for each topology (Figure 2a; Supplementary Table S5). This indicates that many internal branches had lengths close to zero (Züst et al. 2020), giving rise to polytomies that could not be resolved with the 234 single-copy gene sampling used in this analysis.

Although part of these high levels of intragenomic discordance could have been caused by extensive ILS (Stull et al. 2023) in the more recently evolved Loliinae lineages, which likely diverged in Late Pliocene – Quaternary, based on our conservative dating estimates (Figure 4; Minaya et al., 2017; Moreno-Aguilar et al., 2020), they are probably a consequence of the high rates of hybridizations and polyploidizations. Although the conserved and mostly single-copy nuclear gene set of Angiosperms353 has proven to be an invaluable tool for reconstructing the phylogeny of supraspecific plant groups (Baker et al., 2021, 2022), its resolving power at the species and intraspecific levels is less clear. While some studies have demonstrated the ability of these genes to resolve intricate relationships of recently evolved lineages (Thomas et al. 2021), others have emphasized the problems encountered in reconstructing coalescing phylogenies in lineages prone to short- or long-branch attractions and introgressions (Maurin et al. 2021). Even extensive sampling of thousands of nuclear orthologous genes failed to recover quartets support for the species tree in young, highly hybridogenic Brassicaceae groups due to large intragenomic discordance (Züst et al. 2020), or ruled out most of *Brachypodium* gene trees that were topologically incongruent with the diploid species tree (Sancho et al. 2022). Our results are in agreement with these and other works (Philippe et al. 2011; Maurin et al. 2021) which have demonstrated that adding more single-copy genes may not resolve the phylogenies of recently radiated and highly hybridogenic groups. However, a careful gene selection, refinement of analytical processes, and use of concatenation and coalescent approaches (Smith et al. 2020) can help address the problem. Despite the high levels of intragenomic discordance and the low agreement of individual single-copy gene trees with the species trees, the main topologies of the concatenated partitioned ML and the ASTRAL-III species tree (Figures 1, 2a, Supplementary Fig. S3) revealed clades that fitted to taxonomic circumscriptions and/or geographic distributions of most of the studied Loliinae species (Catalán 2006; Catalán et al. 2007a; Minaya et al. 2017; Moreno-Aguilar et al. 2022a).

### Rampant introgressions and allopolyploidizations framed the evolutionary history of the Loliinae

Resolving the phylogeny of Loliinae is challenging, as the subtribe experienced a relatively recent radiation that resulted in a large number of hybridizing species and a prevalence of allopolyploidization (Catalán, 2006; Moreno-Aguilar et al., 2022b). Estimated divergence dates obtained from treePL reveal a main divergence during the mid-Miocene within the crown Loliinae, rendering Broad-leaved and Fine-leaved species. Notably, these results (∼16.02 Ma, Figure 4a) align closely with previous findings (Minaya, 2017; Moreno-Aguilar, 2020). Similarly, the BL clade shows minimal dating variation when compared to the FL clade (16.3 vs 16.06 Ma, Figure 4). However, in this study, we have found differences in age estimates between certain lineages, which could be attributed to the expanded scale of the number of the genes analyzed in our study, together with the integration of data from new species (e. g., Festuca clade, 8.76 – 2.4 Ma; American II clade, 12.37 – 6.73 Ma; Figure 4a). This increased data set contributes to a more complete and robust sample size. In addition, the accelerated radiation of species, with an emphasis on the Fine-leaved clade within the Loliinae, acquires remarkable importance, potentially clarified by the pronounced ploidy diversity and increase in recently evolved lineages (e. g., Festuca, Aulaxyper, American II lineages) (Figures 2a, 2b, 4a, Supplementary Table S1).

The existence of rampant introgressions in Loliinae have been corroborated in our study through the three alternative testing approaches using nuclear-only data and cytonuclear discordance analyses. High levels of hybridization were detected in the ingroup taxa via HyDe tests of the 234 nuclear single copy genes (gamma scores 0.3-0.7; Supplementary Figure S4), while lineage-specific taxa showing signatures of hybridization and polyploidization were recognized through the assessment of high percent values of locus heterozygosity (LH) (> 78%) and allele divergence (AD) (> 1.5%) for the same set of nuclear genes across the studied samples (Figures 2b, 2c; Supplementary Fig. S5). LH vs AD classes tentatively described as old polyploids and high polyploids, encompass putatively hybridogenic taxa within several Loliinae lineages (Figure 2c) although they were predominant in some of them (Supplementary Fig. S5). The incongruence discordance test performed with PACo further confirmed the hybridogenic nature of the 61 mispaired terminals showing strong support in the nuclear and plastid trees (Figures 3, 4; Supplementary Fig. S5). These terminals included lineages or tips with both ancestors evolving within the BL (Subulatae-Hawaiian, Tropical-South African, Mexico-Central-South American) or FL (American II, Aulaxyper, Afroalpine, American-Neozeylandic) clades, and even ‘transclade’ species originating from distantly related BL and FL ancestors (*F. altaica*). In contrast, the most congruent lineages between the two topologies were those of the BL Schedonorus, Subbulbosae, Drymanthele (Phaeochloa) and Lojaconoa, and FL Eskia, Festuca+Wangenheimia and American-Vulpia-Pampas groups (Figures 3, 4; Supplementary Fig. S5). Many of the discordant hybrid terminals correspond to polyploids (Figures 3, 4; Supplementary Table S1; Supplementary Fig. S5), thus reaffirming the pervasiveness of allopolyploidy in the Loliinae, which was also confirmed by the strong asymmetric karyotypes of some of these taxa (Moreno-Aguilar et al., 2022b; Martínez-Segarra, 2021, 2022).

A comparative analysis of nuclear LH – AD classes and cytonuclear PACo discordances across Loliinae lineages revealed an overall correlated trend ranging from non-existent to low-hybridogenesis for diploid and old-polyploid species, topologically congruent in both trees (e. g., BL Drymanthele-Phaeochloa, Lojaconoa, Scariosa, and Pseudoscariosa, FL Psilurus-Vulpia), to high-hybridogenesis for high-polyploid species displaying topological discordances (e. g., BL Coironhuecu, FL Afroalpine, American II, Subulatae-Hawaiian) (Figures 2b, 2c; Supplementary Fig. S5). Accordingly, several Loliinae groups showed concordant but variable LH – AD and PACO patterns for their species, which varied from the non-hybridogenic and congruent features of diploids-tetraploids to the high-hybridogenic and incongruent ones of hexa-to-deca-polyploids (e. g. BL Tropical-South Africa p.p., FL Festuca p.p., Aulaxyper p.p., and the Exaratae-Loretia grade). However, several Loliinae lineages composed exclusively of polyploids presented hybridogenic but topologically congruent taxa (e. g., BL MCSA I group, FL American I, America-Vulpia-Pampas, Fernandezian) (Figures 2b, 2c; Supplementary Fig. S5).

Interestingly, when we examined if levels of hybridogenesis and discordance in the Loliinae could be related to time of divergence or shared ancestry, we did not find any compelling evidence for it (Figures 4a, 4b). By contrast, sister or closely related lineages of similar age had completely different patterns (e.g., congruent American-Vulpia-Pampas and American I (12.58-7.16 Ma) vs. incongruent American II (12.37 Ma) clades). The recently evolved FL Festuca (8.76 Ma) and Aulaxyper (10.68-7.61 Ma) groups also presented contrasted patterns, with high-polyploids of the latter group showing higher LH and AD values and higher discordance than those of the former group (Figures 2b, 2c, 4, S5). Surprisingly, other lineages of similar age, like BL Schedonorus-Lolium (9.62 Ma), that also include a diversity of ploidy-level species (diploids-to-decaploids), exhibited relatively medium to low LH and AD values for its known high-allopolyploids (e.g., *F. arundinacea, F. letoruneuxiana*) and no-topological discordances for any of its taxa (Figures 2a, 2b, 4a, 4b; Supplementary Fig. S5).

These results point towards to evolutionary isolation and hybridogenic nature as the driving factors responsible for the distinct hybridization levels detected in the Loliinae. Taxa of the phylogenetically isolated Schedonorus-Lolium lineage have hybridized extensively among them and mostly between close species (Kopecký and Studer 2014; Glombik et al. 2021), but not or very rarely with taxa from other lineages (Catalán 2006), therefore explaining their high cytonuclear concordance. Within the phylogenetically non-isolated and largest fine-leaved lineages, species of the Festuca clade have created hybrid swarms of increasing ploidy levels but with low intercrossing with less related taxa, thus presenting low cytonuclear discordances. Contrastingly, the highly crossable hexaploid Aulaxyper taxa can easily hybridize with intra- and extra-clade FL Festuca species and other genera (e. g., x *Festulpia*) (Catalán 2006; Catalán et al. 2007), therefore showing high topological discordances (Figures 2a, 2b, 4a, 4b; Supplementary Fig. S5). The abundance of allopolyploid species in the Loliinae phylogeny (70% of samples with known ploidy level; Supplementary Table S1) could have aggravated the levels of discordance. Tracking the evolutionary history of orphan allopolyploids should be accomplished with whole-genome or transcriptome data (Sancho et al. 2022), currently missing for Loliinae. While we have not reconstructed the full evolutionary history of Loliinae, our nuclear single-copy gene and plastome trees have revealed crucial hybridizations patterns in different Loliinae lineages (Figures 2a, 2b, 3, 4; Supplementary Figs. S3, S6).

## Supporting information

Moreno-Aguilar et al. 2024.Suppl.Materials

## Author Contributions

P.C., M.F.M.-A. and J.V. designed the study. M.F.M.-A., J.C.O., G.M.-S., J.A.D., A.S., I.A., W.C. and P.C. collected the samples. M.F.M.-A. developed the experimental work. M.F.M.-A., J.V., A.S.-R., D.C., I.A. and P.C. analyzed the data and interpreted the results. P.C. and M.F.M.-A. wrote the draft manuscript. All authors have read and agreed to the published version of the manuscript.

## Acknowledgements

We thank Nina Probatoba and the AAU, HUTPL, OSC, MO, CONC, US, UZ and VBGI herbaria for lending Loliinae samples for our study, the Ministerio del Ambiente of Ecuador for permitting us to collect Loliinae samples in the Ecuadorian páramos (MAEDNB- CM-2015-0016), and Luis A. Inda for his advice on ploidy levels of Loliinae. Target capture data and genome skimming data of the studied samples were generated at Arbor Biosciences (Ann Arbor, USA) and at the Centro Nacional de Análisis Genómicos (CNAG, Barcelona, Spain) and Macrogen (Madrid, Spain), respectively. The bioinformatic and evolutionary analyses were performed in the Bioflora laboratory of the Escuela Politécnica Superior de Huesca (Universidad de Zaragoza, Spain).

