## Supplementary material for "Genic and phylogenomic discordances reveal conflicting hybridization episodes in temperate Loliinae grasses": Moreno-Aguilar et al. 2024.Suppl.Materials: Moreno-Aguilar et al. 2024_SupplMaterials.docx

**Supplementary materials**

**Supplementary Table S1**. List of taxa included in the single-copy gene phylogenetic study of Loliinae. Sample code (Arbor Biosciences), Taxon (taxon name and authority), Phylogenetic group, Taxonomic classification (subgeneric, sectional), ploidy level, source, and information on genomic data. Gene target capture data (number of Illumina raw reads obtained from gene target capture (raw reads), and Genbank accession code for single-copy-gene data per sample); Genome skimming data (number of million Illumina raw reads obtained from genome skimming (plus insert size), and Genbank accession code for the assembled plastome of each sample). Newly studied species and new genome data are highlighted in bold. Taxonomic classification is based on Alexeev (1977, 1978, 1980, 1981, 1982, 1983, 1984a, 1984b, 1985a, 1985b, 1986); Catalán et al. (2007); Catalán & Muller (2012); Devesa & Martinez-Sagarra (2020); Lu et al. (2006); Moreno-Aguilar et al. (2022); Sainy-Yves (1927); Stančik & Peterson (2007); Tovar (1993); Tzvelev (1971); Tzvelev & Probatova (2019) (see references below). Ploidy levels assigned to each taxon are based on published records.

| **Sample code** | **Taxon** | **Phylogenetic group** | **Taxonomic classification** | **Ploidy** | **Source** | **Gene target capture** | | **Genome skimming** | | |
| --- | --- | --- | --- | --- | --- | --- | --- | --- | --- | --- |
|  |  |  |  |  |  | **No. Illumina raw Reads** | **Genbank accession code** | **No. million of raw reads** | **Insert size** | **Plastome** |
| 1_S1 | *Festuca abyssinica* Hochst. ex A. Rich. | Afroalpine | *incertae sedis* | 4x | Tanzania: Kilimanjaro | 4,614,899 | **SAMN38259822** | 12,041 | 166.00 | SAMN14647043 |
| 2_S112 | *Festuca acuminata* Gaudin | Eskia | *incertae sedis?* (sect. Eskia) | 2x | Switzerland: Neuchatel | 2,795,919 | **SAMN38259823** | 25,027 | 208.00 | **SAMN42486795** |
| 3_S223 | *Festuca africana* (Hack.) Clayton | Tropical-South African | *incertae sedis* | 10x | Uganda: Gahinga | 2,692,853 | **SAMN38259824** | 13,549 | 195.00 | SAMN14647044 |
| 4_S334 | *Festuca aloha* Catalán, Soreng & P.M.Peterson | Subulatae-Hawaiian | *F.* subgen. *Subulatae* (Tzvelev) E. Alexeev | ? | USA: Hawaii | 5,390,793 | **SAMN38259825** | 27,133 | 81.00 | **SAMN42486796** |
| 5_S395 | *Festuca alpina* Suter | Festuca | *F*. subgen. *Festuca* sect. *Festuca* subsect. *Festuca* | 2x | Slovenia: Kammiske Alpe | 1,707,739 | **SAMN38259826** | 19,017 | 201.00 | **SAMN42486797** |
| 6_S406 | ***Festuca altaica* Trin.** | Leucopoa | *F*. subgen. *Leucopoa* (Griseb.) Hack. sect. *Breviaristatae* Krivot. | 4x | Russia: Republic of Altai | 2,346,319 | **SAMN38259827** | 16,817 | 184.50 | **SAMN42486798** |
| 8_S428 | *Festuca amplissima* Rupr. | Ruprechtia - MCSA II | *F*.subgen. *Drymanthele* V.I. Krecz. & Bobrov sect. *Ruprechtia* E.B. Alexeev | 6x | Mexico: Chihuahua | 4,160,251 | **SAMN38259828** | 12,058 | 220.00 | SAMN14647045 |
| 9_S439 | *Festuca andicola* Kunth | American II | *F*.subgen. *Festuca* Sect. *Aulaxyper* Dumort. | 4x | Ecuador: Loja | 2,033,048 | **SAMN38259829** | 10,178 | 212.00 | **SAMN42486799** |
| 10_S2 | *Festuca argentina* (Speg.) Parodi | Coironhuecu | *F*.subgen. *Coironhuecu* Moreno-Aguilar, Arnelas & Catalán | 4x | Argentina: Rio Negro | 2,461,718 | **SAMN38259830** | 22,928 | 213.00 | SAMN30029287 |
| 11_S13 | *Festuca arundinacea atlantigena* (St.-Yves) Auquier | Schedonorus - Lolium | *F.*subgen. *Schenodorus* (P. Beauv.) Peterm. sect. *Schenodorus* | 8x | Morocco: Mahgrebian | 3,444,233 | **SAMN38259831** | 15,091 | 224.00 | SAMN27777775 |
| 12_S24 | *Festuca arundinacea var. letourneuxiana* (St.-Yves) Torrecilla & Catalán | Schedonorus - Lolium | *F.*subgen. *Schenodorus* (P. Beauv.) Peterm. sect. *Schenodorus* | 10x | Morocco:Atlas Mountains | 2,332,917 | **SAMN38259832** | 16,839 | 250.00 | SAMN14647059 |
| 13_S35 | ***Festuca asperula* Vickery** | Australia - Tasmania | *incertae sedis* | ? | Australia: NWS Australia | 4,684,992 | **SAMN38259833** | 12,030 | 122.00 | **PRJNA1073237** |
| 14_S46 | *Festuca asplundii* E.B. Alexeev | American II | *F*.subgen. *Festuca* sect. *Festuca* | 6x | Ecuador: Loja | 2,290,543 | **SAMN38259834** | 25,088 | 300.00 | SAMN14647046 |
| 16_S68 | *Festuca brevipila* R. Tracey | Festuca | *F*.subgen. *Festuca* sect. *Festuca* | 6x | Switzerland: Neuchatel | 3,202,502 | **SAMN38259835** | 12,712 | 190.00 | **SAMN42486800** |
| 17_S79 | *Festuca calabrica* Huter, Porta & Rigo | Leucopoa | *F*.subgen. *Leucopoa* (Griseb.) Hack. sect. *Amphigenes* (Janka)Tzvelev | ? | Italy: Calabria | 2,212,093 | **SAMN38259836** | 9,863 | 248.00 | **SAMN42486801** |
| 18_S90 | *Festuca caldasii* (Kunth) Kunth | Glabricarpae - MCSA I | *F*.subgen. *Subulatae* (Tzvelev) E. Alexeev sect. *Glabricarpae* E.B. Alexeev | 4x | Ecuador: Catamayo | 1,465,186 | **SAMN38259837** | 11,919 | 88.50 | SAMN14647047 |
| 19_S101 | *Festuca californica* Vasey | American - Neozeylandic | *F*.subgen. *Leucopoa* (Griseb.) Hack. sect. *Breviaristatae* Krivot. | 8x | USA: California | 8,643,128 | **SAMN38259838** | 13,430 | 228.00 | **SAMN42486802** |
| 20_S113 | *Festuca capillifolia* Dufour ex Roem. & Schult. | Exaratae - Loretia | *F*.subgen. *Festuca* sect. *Festuca* subsect. *Exaratae* St.-Yves | 2x | Morocco: Middle Atlas | 1,820,969 | **SAMN38259839** | 11,430 | 82.00 | SAMN14647048 |
| 22_S135 | *Festuca caprina* Nees | Afroalpine | *F.*subgen. *Festuca* | 4x | South Africa: Eastern Cape | 3,032,260 | **SAMN38259840** | 29,422 | 150.50 | **SAMN42486803** |
| 21_S124 | ***Festuca carazana* Pilg.** | American II | *incertae sedis* | ? | Peru: Cerro Huiso | 3,716,007 | **SAMN38259841** | 10,937 | 134.00 | **PRJNA1073237** |
| 24_S157 | *Festuca chimborazensis* E.B. Alexeev *subsp. micacochensis* Stančik | American I | *F*.subgen. *Festuca* sect. *Festuca* | 6x | Ecuador: Chimborazo | 1,330,099 | **SAMN38259842** | 10,913 | 254.00 | SAMN14647049 |
| 25_S168 | ***Festuca chodatiana* (St.-Yves) E.B.Alexeev** | Afroalpine | *incertae sedis* | ? | Uganda: Mt. Elgon National Park | 3,950,771 | **SAMN38259843** | 10,362 | 155.50 | **PRJNA1073237** |
| 26_S179 | *Festuca coerulescens* Desf. | Lojaconoa | *F*.subgen. *Festuca* sect. *Lojaconoa* Catalán & Joch. Müll | 2x | Morocco: Rif Mountains | 4,145,191 | **SAMN38259844** | 12,481 | 188.50 | **SAMN39823839** |
| 27_S190 | ***Festuca compressifolia* Presl.** | American II | *incertae sedis* | ? | Peru: Huancavelica | 4,409,347 | **SAMN38259845** | 11,308 | 163.00 | **SAMN42486804** |
| 28_S201 | *Festuca costata* Nees | Tropical - South African | *incertae sedis* | 4x | South Africa: Eastern Cape | 3,843,435 | **SAMN38259846** | 14,693 | 137.00 | **PRJNA1073237** |
| 29_S212 | ***Festuca dasyantha* Kunth** | American II | *F*.subgen. *Festuca* sect. *Cataphyllophorae* E.B. Alexeev | ? | Ecuador: Imbabura | 4,092,718 | **SAMN38259847** | 14,204 | 109.00 | **SAMN42486805** |
| 30_S224 | ***Festuca dichoclada* Pilg.** | Erosiflorae - MCSA I | *F*.subgen. *Erosiflorae* E.B. Alexeev | ? | Peru: Cuzco | 4,078,082 | **SAMN38259848** | 12,466 | 181.00 | SAMN30029291 |
| 31_S235 | ***Festuca distichovaginata* Pilg.** | American II | *incertae sedis* | ? | Peru: Puna | 3,898,537 | **SAMN38259849** | 16,098 | 118.50 | **SAMN42486806** |
| 32_S246 | ***Festuca dolichophylla* J.Presl** | American II | *incertae sedis* | 6x | Peru: Lima | 5,142,446 | **SAMN38259850** | 11,952 | 133.00 | **PRJNA1073237** |
| 33_S257 | *Festuca dracomontana* H.P. Linder | Schedonorus - Lolium | *incertae sedis?(subgen. Schedonorus?)* | ? | South Africa: TVL | 3,601,332 | **SAMN38259851** | 15,835 | 139.00 | SAMN27777776 |
| 34_S268 | *Festuca drymeja* Mert. & W.D.J.Koch | Drymanthele- Phaeochloa | *F*.subgen. *Drymanthele* V.I. Krecz. & Bobrov sect. *Phaeochloa* Griseb. | 2x | Russia: Azerbaidzhancoll | 3,426,473 | **SAMN38259852** | 16,225 | 82.25 | **SAMN39823840** |
| 35_S279 | *Festuca durandoi* Clauson | Subbulbosae | *F*.subgen. *Festuca* sect. *Subbulbosae* Nyman. ex Hack. | 2x | Portugal: Serra Arga | 3,781,121 | **SAMN38259853** | 12,688 | 217.00 | SAMN14647050 |
| 36_S290 | *Festuca elegans* Boiss. | Eskia | *F*.subgen. *Festuca* sect. *Pseudatropis* Krivot. | 4x | Spain: CC03 | 3,071,001 | **SAMN38259854** | 10,765 | 182.00 | **PRJNA1073237** |
| 37_S301 | *Festuca engleri* Pilg. | Tropical - South African | *incertae sedis* (*Pseudobromus* *engleri* (Pilg.) Clayton) | ? | Kenya: Mt. Kenya | 3,012,262 | **SAMN38259855** | 16,534 | 115.50 | **SAMN42486807** |
| 39_S323 | *Festuca extremiorientalis* Owhi | Subulatae - Hawaiian | *F*.subgen. *Subulatae* (Tzvelev) E. Alexeev sect. *Subulatae* | 4x | Japan: Tohoku | 5,875,021 | **SAMN38259856** | 15,252 | 118.00 | **PRJNA1073237** |
| 40_S335 | *Festuca fenas* Lag | Schedonorus - Lolium | *F*.subgen. *Schenodorus* (P. Beauv.) Peterm. sect. *Schenodorus* | 4x | Spain:W Mediterranean | 2,660,548 | **SAMN38259857** | 16,112 | 271.00 | SAMN14647052 |
| 41_S346 | *Festuca filiformis* Pourr. | Festuca | *F*.subgen. *Festuca* sect. *Festuca* | 2x | Switzerland: Neuchatel | 2,372,826 | **SAMN38259858** | 10,768 | 222.00 | **PRJNA1073237** |
| 43_S368 | *Festuca flacca* Hack. ex E.B.Alexeev | American II | *F*.subgen. *Subulatae* (Tzvelev) E. Alexeev sect. *Subulatae* | 4x | Ecuador: Pichincha | 1,950,954 | **SAMN38259859** | 14,188 | 170.25 | **PRJNA1073237** |
| 44_S379 | *Festuca fontqueri* St.-Yves | Schedonorus - Lolium | *F*.subgen. *Schenodorus* (P. Beauv.) Peterm. sect. *Schenodorus* | 2x | Morocco: Rif Mountains | 2,686,917 | **SAMN38259860** | 22,187 | 300.00 | SAMN14647054 |
| 45_S390 | *Festuca francoi* Fern. Prieto, C. Aguiar, E. Días & M.I. Gut | Aulaxyper | *F*.subgen. *Festuca* sect. *Aulaxyper* Dumort. | 2x | Portugal: Açores | 1,923,931 | **SAMN38259861** | 17,592 | 186.00 | SAMN14647057 |
| 46_S391 | *Festuca gautieri* (Hack.) K.Richt. | Eskia | *F*.subgen. *Festuca* sect. *Eskia* Willk. | 2x | Spain:Granada | 3,420,343 | **SAMN38259862** | 13,941 | 177.00 | SAMN30029292 |
| 47_S392 | *Festuca gigantea* (L.) Vill. | Schedonorus - Lolium | *F*.subgen. *Schenodorus* (P. Beauv.) Peterm. sect. *Plantynia* (Dumort.) Tzvelev | 6x | Norway | 2,088,065 | **SAMN38259863** | 20,914 | 223.00 | SAMN27777777 |
| 48_S393 | *Festuca glauca* Vill. | Festuca | *F.*subgen. *Festuca* sect. *Festuca* | 2x | Spain: Barcelona | 1,783,800 | **SAMN38259864** | 35,888 | 200.00 | **PRJNA1073237** |
| 49_S394 | ***Festuca glumosa* Hack. ex E.B.Alexeev** | American II | *F*.subgen. *Festuca* sect. *Festuca* | 4x | Ecuador: Cotopaxi | 2,170,353 | **SAMN38259865** | 15,202 | 209.00 | **SAMN42486808** |
| 50_S396 | *Festuca gracilior* (Hack.) Markgr.-Dann | Festuca | *F*.subgen. *Festuca* sect. *Festuca* | 2x | Spain: Barcelona | 2,593,769 | **SAMN38259866** | 13,184 | 199.50 | **PRJNA1073237** |
| 51_S397 | *Festuca gracillima* Hook. F. | American - Neozeylandic | *incertae sedis* | 6x | Argentina: Tierra de Fuego | 3,291,849 | **SAMN38259867** | 13,888 | 224.00 | SAMN14647055 |
| 52_S398 | *Festuca gudoschnikovii* Stepanov | Schedonorus - Lolium | *F*.subgen. *Schenodorus* (P. Beauv.) Peterm. sect. *Plantynia* (Dumort.) Tzvelev? | 4x | Russia: Krasnoyarskii Krai | 2,795,488 | **SAMN38259868** | 13,994 | 208.75 | SAMN27777778 |
| 53_S399 | *Festuca henriquesii* Hack. | Aulaxyper | *F.*subgen. *Festuca* sect. *Aulaxyper* Dumort.?/Festuca? | 2x | Portugal: Torre | 2,560,289 | **SAMN38259869** | 12,574 | 185.00 | **PRJNA1073237** |
| 54_S400 | *Festuca hephaestophilla* (Nees) Nees. | Exaratae - Loretia | *F*.subgen. *Festuca* | 4x | Mexico: Nuevo Leon | 3,189,816 | **SAMN38259870** | 10,690 | 150.00 | **SAMN42486809** |
| 55_S401 | *Festuca hieronymi* Hack. | American II | *F*.subgen. *Festuca* | 6x | Argentina: Cordoba | 4,080,374 | **SAMN38259871** | 14,242 | 177.50 | **SAMN42486810** |
| 56_S402 | *Festuca holubii* Stančík | American I | *F*.subgen. *Festuca* sect. *Festuca* | ? | Ecuador: Saraguro | 2,014,431 | **SAMN38259872** | 10,264 | 249.00 | SAMN14647056 |
| 57_S403 | ***Festuca humilior* Nees** | American II | *incertae sedis* | ? | Peru: Tarma | 2,883,239 | **SAMN38259873** | 14,100 | 67.00 | **SAMN42486811** |
| 58_S404 | *Festuca hystrix* Boiss. | Festuca | *F*.subgen. *Festuca* sect. *Festuca* | 2x | Spain: Almeria | 3,277,551 | **SAMN38259874** | 13,029 | 188.00 | **PRJNA1073237** |
| 59_S405 | *Festuca iberica* (Hack.) K.Richt. | Aulaxyper | *F*.subgen. *Festuca* sect. *Aulaxyper* Dumort. | 6x | Spain: Granada | 3,816,394 | **SAMN38259875** | 11,016 | 155.00 | **SAMN39823841** |
| 60_S407 | *Festuca imbaburensis* Stančik | American I | *F*.subgen. *Festuca* sect. *Festuca* | 4x | Ecuador: Chimborazo | 1,780,378 | **SAMN38259876** | 19,836 | 211.00 | **PRJNA1073237** |
| 61_S408 | *Festuca indigesta* Boiss. | Festuca | *F*.subgen. *Festuca* sect. *Festuca* | 6x | Spain: Almería | 1,892,280 | **SAMN38259877** | 15,233 | 194.00 | **SAMN42486812** |
| 63_S410 | ***Festuca kolesnikovii* Tzvelev** | Festuca | *incertae sedis* | ? | Russia: Primorskii Krai | 3,561,231 | **SAMN38259878** | 18,310 | 105.00 | **SAMN42486813** |
| 64_S411 | ***Festuca kurtziana* St.-Yves** | American - Neozeylandic | *F*.subgen. *Festuca* sect. *Festuca* | 6x | Argentina: Mendoza | 4,043,339 | **SAMN38259879** | 8,555 | 200.00 | **PRJNA1073237** |
| 65_S412 | ***Festuca laegaardii* Stančik** | American II | *F*.subgen. *Festuca* sect *Cataphyllophorae* E.B. Alexeev | 4x | Ecuador: Azuay | 2,064,736 | **SAMN38259880** | 15,489 | 217.00 | **SAMN42486814** |
| 66_S413 | *Festuca laevigata* Gaudin | Festuca | *F*.subgen. *Festuca* sect. *Festuca* | 8x | Spain: Cataluña | 2,811,015 | **SAMN38259881** | 15,693 | 198.00 | **SAMN42486815** |
| 67_S414 | *Festuca lasto* Boiss*.* | Drymanthele - Phaeochloa | *F*.subgen. *Drymanthele* sect. *Phaeochloa* Griseb. | 2x | Spain: Cadiz | 2,353,759 | **SAMN38259882** | 21,581 | 300.00 | SAMN14647058 |
| 68_S415 | *Festuca lemanii* Bastard | Festuca | *F*.subgen. *Festuca* sect. *Festuca* | 6x | Spain: Girona | 1,864,653 | **SAMN38259883** | 13,309 | 209.50 | **SAMN42486816** |
| 69_S416 | ***Festuca leptopogon* Stapf** | Subulatae - Hawaiian | *F*.subgen. *Subulatae* (Tzvelev) E. Alexeev sect. *Subulatae* | 4x | China: Xizang | 1,472,141 | **SAMN38259884** | ----- | ----- | ----- |
| 70_S418 | *Festuca longiauriculata* Fuente, Ortúñez & Ferrero Lom. | Festuca | *F*.subgen. *Festuca* sect. *Festuca* | 2x | Spain: Almería | 2,792,617 | **SAMN38259885** | 15,218 | 200.00 | **PRJNA1073237** |
| 71_S419 | ***Festuca longipes* Stafp** | Tropical - South African | *incertae sedis* | ? | South Africa: SA 031 E Cape | 1,637,913 | **SAMN38259886** | 15,404 | 169.00 | **PRJNA1073237** |
| 72_S420 | ***Festuca lugens* (E.Fourn.) Hitchc. ex Hern.-Xol.** | Asperifolia - MCSA I | *F*.subgen. *Asperifolia* E.B. Alexeev | 4x | Honduras: Morazán | 2,558,095 | **SAMN38259887** | ----- | ----- | ----- |
| 73_S421 | *Festuca mairei* St.-Yves | Schedonorus - Lolium | *F*.subgen. *Festuca* sect. *Scariosae* Hack. | 4x | Morocco: Atlas Mountains | 4,423,842 | **SAMN38259888** | 19,134 | 209.00 | SAMN14647060 |
| 74_S422 | *Festuca marginata* (Hack.) K. Richt. | Festuca | *F*.subgen. *Festuca* sect. *Festuca* | 2x | Spain: Montsec | 1,980,381 | **SAMN38259889** | 14,637 | 254.00 | **SAMN42486817** |
| 75_S423 | *Festuca mekiste* Clayton | Tropical - South African | *incertae sedis* | ? | Kenya: Mt. Elgon National Park | 2,570,539 | **SAMN38259890** | 16,245 | 201.00 | SAMN27777779 |
| 76_S424 | *Festuca modesta* Steud. | Drymanthele - Muticae | *F*.subgen. *Drymanthele* V.I. Krecz. & Bobrov sect. *Muticae* S.L. Lu | 2x | India: NW Himalaya | 3,735,505 | **SAMN38259891** | ----- | ----- | ----- |
| 77_S425 | ***Festuca mollissima* V.I. Krecz. & Bobrov** | Festuca | *incertae sedis* | 2x | Russia: Primorskii Krai | 4,383,423 | **SAMN38259892** | 13,510 | 80.50 | **SAMN42486818** |
| 78_S426 | *Festuca molokaiensis* Soreng, P.M. Peterson & Catalán | Subulatae - Hawaiian | *incertae sedis* | ? | USA: Hawai | 2,703,379 | **SAMN38259893** | 12,188 | 100.00 | SAMN14647061 |
| 79_S427 | ***Festuca monguensis* Stančik** | American II | *F*.subgen. *Festuca* sect. *Festuca* | ? | Colombia: Boyacá | 6,603,137 | **SAMN38259894** | 13,545 | 219.00 | **PRJNA1073237** |
| 80_S429 | ***Festuca muelleri* Vickery** | Leucopoa | *incertae sedis* | ? | Australia: Australia Capital Territory | 3,806,324 | **SAMN38259895** | 14,886 | 69.50 | **SAMN42486819** |
| 81_S430 | *Festuca nevadensis* (Hack.) K. Richt. | Aulaxyper | *F*.subgen. *Festuca* Sect. *Aulaxyper* Dumort. | 10x | Spain: Almeria | 6,011,636 | **SAMN38259896** | 13,223 | 95.50 | **SAMN42486820** |
| 82_S431 | *Festuca nigrescens* Lam. | Aulaxyper | *F*.subgen. *Festuca* sect. *Aulaxyper* Dumort. | 6x | Switzerland: Neuchatel | 2,205,457 | **SAMN38259897** | 23,337 | 175.00 | **PRJNA1073237** |
| 83_S432 | ***Festuca olgae* (Regel) Krivot.** | Leucopoa | *F*.subgen. *Leucopoa* (Griseb.) Hack. | ? | Kazakhstan: West Tyan-Shan mts. | 2,771,621 | **SAMN38259898** | 18,527 | 300.00 | **PRJNA1073237** |
| 84_S433 | *Festuca orthophylla* Pilg. | American II | *F*.subgen. *Festuca* | 8x | Argentina: Jujuy | 3,615,539 | **SAMN38259899** | 13,484 | 38.00 | **PRJNA1073237** |
| 85_S434 | *Festuca ovina* L. | Festuca | *F*.subgen. *Festuca* sect. *Festuca* | 2x | Germany: Thüringen | 3,117,984 | **SAMN38259900** | 11,364 | 188.00 | SAMN14647062 |
| 86_S435 | *Festuca pampeana* Speg. | American - Vulpia - Pampas | *F.*subgen. *Festuca* | 8x | Argentina: Buenos Aires | 2,867,938 | **SAMN38259901** | 14,862 | 215.00 | SAMN14647063 |
| 87_S436 | *Festuca paniculata* (L.) Schinz & Thell | Subbulbosae | *F*.subgen. *Festuca* sect. *Subbulbosae* Nyman ex Hack. | 2x | Spain: Caceres | 5,807,680 | **SAMN38259902** | 35,808 | 223.00 | SAMN14647064 |
| 88_S437 | ***Festuca parciflora* Swallen** | American II | *F*.subgen. *Festuca* sect. *Festuca* | 4x | Ecuador: Cotopaxi | 2,315,801 | **SAMN38259903** | 16,413 | 300.00 | **PRJNA1073237** |
| 89_S438 | *Festuca parvigluma* Steud. | Subulatae - Hawaiian | *F*.subgen. *Subulatae* sect. *Subulatae* (Tzvelev) E. Alexeev | 4x | China: Baotianman | 4,354,758 | **SAMN38259904** | 15,872 | 223.00 | SAMN14647065 |
| 90_S440 | *Festuca pilgeri* St.-Yves | Afroalpine | *F*.subgen. *Festuca* | ? | Kenya | 4,257,594 | **SAMN38259905** | 20,003 | 190.00 | **SAMN39823842** |
| 91_S441 | *Festuca plebeia* R.Br. | Australia - Tasmania | *incertae sedis* | ? | Australia: Proctors Road | 2,646,006 | **SAMN38259906** | 15,479 | 165.00 | **SAMN42486821** |
| 92_S442 | *Festuca plicata* Hack. | Exaratae - Loretia | *F*.subgen. *Festuca* sect. *Festuca* | 2x | Spain: Granada | 2,499,030 | **SAMN38259907** | 14,451 | 81.50 | **SAMN42486822** |
| 93_S443 | *Festuca pratensis* Huds. | Schedonorus - Lolium | *F.*subgen*. Schenodorus* (P. Beauv.) Peterm. sect. *Schenodorus* | 2x | United Kingdom: England | 2,431,565 | **SAMN38259908** | 30,021 | 182.50 | SAMN14647066 |
| 94_S444 | *Festuca procera* Kunth | American II | *F*.subgen. *Festuca* sect. *Cataphyllophorae* E.B. Alexeev | 4x | Ecuador: Chimborazo | 2,280,684 | **SAMN38259909** | 40.669 | 271.00 | SAMN14647067 |
| 95_S445 | *Festuca pseudeskia* Boiss. | Pseudoscariosa | *F*.subgen. *Festuca* sect. *Pseudoscariosa* Krivot | 2x | Spain: Granada | 2,551,426 | **SAMN38259910** | 14,767 | 263.00 | **PRJNA1073237** |
| 96_S446 | *Festuca pumila* Wilk. | Eskia | *F*.subgen. *Festuca* sect. *Eskia* Willk. | 2x | Austria: Steiermark | 3,045,899 | **SAMN38259911** | 19,264 | 170.00 | **PRJNA1073237** |
| 97_S447 | *Festuca pyrenaica* Reut. | Exaratae - Loretia | *F*.subgen. *Festuca* sect. *Festuca* subsect. *Exaratae* St.-Yves? | 4x | Spain: Huesca | 1,298,678 | **SAMN38259912** | 30.021 | 224.50 | SAMN14647068 |
| 98_S448 | *Festuca pyrogea* Speg. | Festuca | *F*.subgen. *Festuca* sect. *Festuca* subsect. *Festuca* | ? | Argentina: Tierra de fuego | 2,921,816 | **SAMN38259913** | 16,835 | 300.00 | SAMN14647069 |
| 99_S449 | *Festuca quadridentata* Kunth | Erosiflorae - MCSA II | *F*.subgen. *Erosiflorae* E.B. Alexeev | ? | Ecuador: Chimborazo | 3,161,560 | **SAMN38259914** | 15,091 | 193.00 | SAMN14647070 |
| 100_S3 | ***Festuca queriana* Litard.** | Aulaxyper | *F*.subgen. *Festuca* Sect. *Aulaxyper* Dumort. | 4x | Spain: Zamora | 1,837,449 | **SAMN38259915** | 18,691 | 130.00 | **SAMN42486823** |
| 101_S4 | *Festuca raddei* Enustsch. & Prob*. [sp. nova (F. aggr. rubra)]* | Aulaxyper | *incertae sedis* | ? | Russia: Far East | 1,501,208 | **SAMN38259916** | 14,121 | 195.50 | **SAMN42486824** |
| 102_S5 | *Festuca reverchonii* Hack. | Festuca | *F*.subgen. *Festuca* sect. *Festuca* | 2x | Spain: Jaén | 1,302,842 | **SAMN38259917** | 14,565 | 141.50 | **SAMN42486825** |
| 103_S6 | *Festuca richardsonii* Hook. | Aulaxyper | *incertae sedis? subgen. Festuca sect. Aulaxyper?* | 6x | Uganda: Chukotskii Natsional’nyi Okrug | 2,005,109 | **SAMN38259918** | 14,041 | 169.50 | **PRJNA1073237** |
| 104_S7 | ***Festuca rigidifolia* Tovar** | American II | *incertae sedis* | ? | Peru: Lima | 2,440,444 | **SAMN38259919** | 14,379 | 134.50 | **SAMN42486826** |
| 105_S8 | *Festuca rubra* L. | Aulaxyper | *F*.subgen. *Festuca* Sect. *Aulaxyper* Dumort. | 6x | Argentina: Tierra de fuego | 1,379,087 | **SAMN38259920** | 25,260 | 100.50 | SAMN27777780 |
| 106_S9 | ***Festuca samensis* Joch.Müll.** | American - Vulpia - Pampas | *F*.subgen. *Festuca* | ? | Bolivia: Santa Cruz | 3,238,719 | **SAMN38259921** | 18,985 | 220.00 | **SAMN42486827** |
| 107_S10 | *Festuca scabra* Vahl | Tropical - South African | *incertae sedis* | 4x | South Africa: KwaZulu Natal | 3,132,091 | **SAMN38259922** | 21,174 | 66.5 | SAMN27777781 |
| 108_S11 | *Festuca scariosa* Pau | Scariosae | *F*.subgen. *Festuca* sect. *Scariosae* Hack. | 2x | Spain: Córdoba | 4,343,876 | **SAMN38259923** | 16,170 | 211.00 | **SAMN42486828** |
| 109_S12 | ***Festuca setifolia* Steud. ex Griseb.** | American II | *F*.subgen. *Festuca* | ? | Peru:Ancash | 3,192,887 | **SAMN38259924** | 19,074 | 153.5 | **PRJNA1073237** |
| 111_S15 | *Festuca simensis* Hochst. ex A.Rich. | Schedonorus - Lolium | *F*.subgen. *Schenodorus* (P. Beauv.) Peterm. sect. *Schenodorus* | 4x | Kenya: Mt. Kenya | 2,097,137 | **SAMN38259925** | 14,159 | 149.0 | SAMN27777782 |
| 112_S16 | ***Festuca sodiroana* Hack. ex E.B. Alexeev** | American II | *F*.subgen. *Subulatae* (Tzvelev) E. Alexeev sect. *Subulatae* | 4x | Ecuador: Azuay | 2,220,817 | **SAMN38259926** | 17,431 | 192.5 | **PRJNA1073237** |
| 114_S18 | *Festuca spectabilis* Jan. | Leucopoa | *F*.subgen. *Leucopoa* (Griseb.) Hack. sect. *Leucopoa* | 6x | Bosnia-Hercegovina: Troglav | 2,399,570 | **SAMN38259927** | 12,960 | 221.00 | SAMN14647071 |
| 115_S19 | *Festuca subuliflora* Scribn. | Subuliflorae | *F*.subgen. *Subuliflorae* E.B. Alexeev sect. *Subuliflorae* | 4x | USA: Marion county | 2,500,554 | **SAMN38259928** | 18,189 | 209.5 | **PRJNA1073237** |
| 116_S20 | *Festuca subulifolia* Benth. | American II | *F*.subgen. *Festuca* sect. *Festuca* | 4x | Ecuador: Chimborazo | 2,742,422 | **SAMN38259929** | 20,276 | 79.0 | **PRJNA1073237** |
| 117_S21 | *Festuca subverticillata* (Pers.) E.B.Alexeev | American II | *F*.subgen. *Leucopoa* (Griseb.) Hack. sect. *Obtusae* (E.B. Alexeev) E.B. Alexeev | 4x | USA: Missouri | 3,251,197 | **SAMN38259930** | 19,399 | 214.0 | **PRJNA1073237** |
| 118_S22 | *Festuca superba* Parodi ex Türpe | Drymanthele s. l. - MCSA I | *F*.subgen. *Drymanthele* V.I. Krecz. & Bobrov | 8x | Argentina: Jujuy | 1,746,583 | **SAMN38259931** | 12,193 | 82.0 | SAMN14647072 |
| 119_S23 | *Festuca triflora* J.F. Gmel. | Lojaconoa | *F*.subgen. *Festuca* sect. *Lojaconoa* Catalán & Joch. Müll | 2x | Morocco: Rif Mountains | 3,566,291 | **SAMN38259932** | 24,472 | 300.00 | SAMN14647073 |
| 23_S146 | *Festuca valdesii* Gonz.-Led. & S.D.Koch | Ruprechtia - MCSA II | *F*.subgen. *Drymanthele* V.I. Krecz. & Bobrov sect. *Ruprechtia* E.B. Alexeev | ? | Mexico: Coahuila | 4,088,715 | **SAMN38259933** | 10.937 | 200.0 | SAMN30029295 |
| 122_S27 | ***Festuca versuta* Beal** | American II | *F*.subgen. *Drymanthele* V.I. Krecz. & Bobrov sect. *Texanae* E.B. Alexeev | ? | USA: Texas | 3,378,862 | **SAMN38259934** | 3,938 | 175.0 | **PRJNA1073237** |
| 123_S28 | *Festuca viviparoidea* Krajina ex Pavlick | Festuca | *F*.subgen. *Festuca* sect. *Festuca* subsect. *Festuca* | 8x | Canada: Northwest territories | 3,549,531 | **SAMN38259935** | 18,988 | 0.00 | **PRJNA1073237** |
| 124_S29 | ***Festuca weberbaueri* Pilg.** | American II | *F*.subgen. *Festuca* | ? | Peru: Recuay | 3,962,222 | **SAMN38259936** | 16,083 | 157.0 | **SAMN42486829** |
| 125_S30 | ***Festuca woronowii* Hack.** | Eskia | *F*.subgen. *Festuca* sect. *Eskia* Willk. | 2x | Russia: Dagestan | 2,901,261 | **SAMN38259937** | 14,067 | 132.5 | **PRJNA1073237** |
| 126_S31 | *Festuca yalaensis* Joch.Müll. & Catalán | American II | *incertae sedis* | ? | Argentina: Jujuy | 3,283,076 | **SAMN38259938** | 18,550 | 115.5 | **SAMN42486830** |
| 127_S32 | *Festuca yvesii* Sennen & Pau | Festuca | *F*.subgen. *Festuca* sect. *Festuca* | 8x | Spain: Girona | 2,914,782 | **SAMN38259939** | 19,772 | 162.0 | **PRJNA1073237** |
| 128_S33 | *Hellerochloa livida* (Kunt) Willd. ex Spreng. | American II | *Hellerochloa* Rauschert | ? | Mexico:Veracruz | 2,573,854 | **SAMN38259940** | 1,563 | 213.5 | **PRJNA1073237** |
| 129_S34 | *Lolium canariense* Steud | Schedonorus - Lolium | *Lolium* L. | 2x | Spain: Canary islands | 1,837,344 | **SAMN38259941** | 16,359 | 231.5 | SAMN27777783 |
| 130_S36 | *Lolium perenne* L. | Schedonorus - Lolium | *Lolium* L. | 2x | United Kingdom: Wales | 2,387,236 | **SAMN38259942** | 28,103 | 244.00 | SAMN27777784 |
| 132_S38 | *Lolium rigidum* Gaudin | Schedonorus - Lolium | *Lolium* L. | 2x | Turkey: USDA | 2,693,803 | **SAMN38259943** | 16,730 | 201.5 | SAMN27777786 |
| 133_S39 | *Lolium saxatile* H. Scholz & S. Scholz | Schedonorus - Lolium | *Lolium* L. | 2x | Spain: Canary islands | 2,910,153 | **SAMN38259944** | 16,001 | 236.0 | SAMN27777787 |
| 134_S40 | *Lolium temulentum* L. | Schedonorus - Lolium | *Lolium* L. | 2x | Turkey | 2,273,535 | **SAMN38259945** | ----- | ----- | ----- |
| 135_S41 | *Megalachne berteroniana* Steud. | Fernandezian | *Megalachne* Steud. | ? | Chile: Juan Fernandez archipelago | 2,594,653 | **SAMN38259946** | 5,288 | 184.00 | SAMN14647074 |
| 136_S42 | *Micropyropsis tuberosa* Romero-Zarco & Cabezudo | Schedonorus - Lolium | *Micropyropsis* Romero Zarco & Cabezudo | 2x | Spain: Huelva | 3,692,416 | **SAMN38259947** | 19,803 | NA | SAMN27777788 |
| 137_S43 | *Psilurus incurvus* (Gouan) Schinz & Thell. | Psilurus - Vulpia | *Psilurus* Trin. | 4x | Spain: Caceres | 3,620,406 | **SAMN38259948** | 15,885 | NA | **SAMN39823843** |
| 138_S44 | *Vulpia ciliata* Dumort. | Psilurus - Vulpia | *Vulpia C.C. Gmel. sect. Vulpia* | 4x | Spain: Toledo | 5,087,200 | **SAMN38259950** | 11,801 | 155.0 | SAMN14647076 |
| 140_S47 | *Vulpia membranacea* (L.) Dumort. | Exaratae - Loretia | *Vulpia* sect. C.C. Gmel. *Monachne* Dumort. | 2x | Spain: Huelva | 2,112,044 | **SAMN38259951** | 21775 | 220.00 | **SAMN42486831** |
| 142_S49 | *Vulpia muralis* (Kunth) Nees | Aulaxyper | *Vulpia* C.C. Gmel. sect. *Vulpia* | 2x | Spain:Caceres | 2,615,655 | **SAMN38259949** | 27,825 | 124.5 | **SAMN42486832** |
| 143_S50 | *Vulpia sicula* (C. Presl) Link | Exaratae - Loretia | *Vulpia* C.C. Gmel. sect. *Loretia* (Duval-Jouve) Boiss. | 2x | Italy: Sicilia | 3,592,248 | **SAMN38259952** | 11,327 | 223.00 | SAMN14647077 |
| 144_S51 | *Wangenheimia lima* (L.) Trin. | Festuca | *Wangenheimia* Moench | 2x | Spain: Zaragoza | 3,318,538 | **SAMN38259953** | 24,316 | 117.0 | **SAMN42486833** |

**Supplementary Table S2**. Summary statistics from HybPiper analysis, based on target gene capture in 132 Loliinae taxa for 234 filtered genes used in this study. Taxon (name and authority), sample code (Arbor Biosciences), number of genes per taxon, percentage of total number of genes per taxon, sequence length of genes (bp) per taxon, percentage of sequence length of genes (bp) per taxon.

| **Specie** | **Sample code** | **No. genes/ taxon** | **% genes/ taxon** | **length of genes/ taxon** | **%length of genes/ taxon** |
| --- | --- | --- | --- | --- | --- |
| *Festuca abyssinica* Hochst. ex A. Rich. | 1_S1 | 212 | 90.60 | 120555 | 66.62 |
| *Festuca acuminata* Gaudin | 2_S112 | 218 | 93.16 | 147978 | 81.78 |
| *Festuca africana* (Hack.) Clayton | 3_S223 | 162 | 69.23 | 76851 | 42.47 |
| *Festuca aloha* Catalán, Soreng & P.M.Peterson | 4_S334 | 214 | 91.45 | 100344 | 55.46 |
| *Festuca alpina* Suter | 5_S395 | 215 | 91.88 | 128505 | 71.02 |
| *Festuca altaica* Trin. | 6_S406 | 221 | 94.44 | 141048 | 77.95 |
| *Festuca amplissima* Rupr. | 8_S428 | 214 | 91.45 | 122259 | 67.57 |
| *Festuca andicola* Kunth | 9_S439 | 212 | 90.60 | 127656 | 70.55 |
| *Festuca argentina* (Speg.) Parodi | 10_S2 | 217 | 92.74 | 134412 | 74.28 |
| *Festuca arundinacea atlantigena* (St.-Yves) Auquier | 11_S13 | 212 | 90.60 | 128154 | 70.82 |
| *Festuca arundinacea var. letourneuxiana* (St.-Yves) Torrecilla & Catalán | 12_S24 | 216 | 92.31 | 127359 | 70.39 |
| *Festuca asperula* Vickery | 13_S35 | 221 | 94.44 | 134349 | 74.25 |
| *Festuca asplundii* E.B. Alexeev | 14_S46 | 221 | 94.44 | 131190 | 72.50 |
| *Festuca brevipila* R. Tracey | 16_S68 | 182 | 77.78 | 90711 | 50.13 |
| *Festuca calabrica* Huter, Porta & Rigo | 17_S79 | 218 | 93.16 | 135465 | 74.86 |
| *Festuca caldasii* (Kunth) Kunth | 18_S90 | 200 | 85.47 | 114345 | 63.19 |
| *Festuca californica* Vasey | 19_S101 | 213 | 91.03 | 105684 | 58.41 |
| *Festuca capillifolia* Dufour ex Roem. & Schult. | 20_S113 | 219 | 93.59 | 141708 | 78.32 |
| *Festuca carazana* Pilg. | 21_S124 | 226 | 96.58 | 119826 | 66.22 |
| *Festuca caprina* Nees | 22_S135 | 219 | 93.59 | 131508 | 72.68 |
| *Festuca valdesii* Gonz.-Led. & S.D.Koch | 23_S146 | 217 | 92.74 | 119376 | 65.97 |
| *Festuca chimborazensis* E.B. Alexeev *subsp. micacochensis* Stančik | 24_S157 | 208 | 88.89 | 127212 | 70.30 |
| *Festuca chodatiana* (St.-Yves) E.B.Alexeev | 25_S168 | 219 | 93.59 | 123924 | 68.49 |
| *Festuca coerulescens* Desf. | 26_S179 | 221 | 94.44 | 145095 | 80.19 |
| *Festuca compressifolia* Presl. | 27_S190 | 203 | 86.75 | 102822 | 56.82 |
| *Festuca costata* Nees | 28_S201 | 221 | 94.44 | 139053 | 76.85 |
| *Festuca dasyantha* Kunth | 29_S212 | 206 | 88.03 | 100920 | 55.77 |
| *Festuca dichoclada* Pilg. | 30_S224 | 226 | 96.58 | 126864 | 70.11 |
| *Festuca distichovaginata* Pilg. | 31_S235 | 208 | 88.89 | 106326 | 58.76 |
| *Festuca dolichophylla* J.Presl | 32_S246 | 225 | 96.15 | 127797 | 70.63 |
| *Festuca dracomontana* H.P. Linder | 33_S257 | 214 | 91.45 | 105360 | 58.23 |
| *Festuca drymeja* Mert. & W.D.J.Koch | 34_S268 | 215 | 91.88 | 119313 | 65.94 |
| *Festuca durandoi* Clauson | 35_S279 | 203 | 86.75 | 119835 | 66.23 |
| *Festuca elegans* Boiss. | 36_S290 | 211 | 90.17 | 110739 | 61.20 |
| *Festuca engleri* Pilg. | 37_S301 | 216 | 92.31 | 121266 | 67.02 |
| *Festuca extremiorientalis* Owhi | 39_S323 | 223 | 95.30 | 119316 | 65.94 |
| *Festuca fenas* Lag*.* | 40_S335 | 210 | 89.74 | 120066 | 66.35 |
| *Festuca filiformis* Pourr. | 41_S346 | 223 | 95.30 | 139326 | 77.00 |
| *Festuca flacca* Hack. ex E.B.Alexeev | 43_S368 | 196 | 83.76 | 92085 | 50.89 |
| *Festuca fontqueri* St.-Yves | 44_S379 | 219 | 93.59 | 144963 | 80.11 |
| *Festuca francoi* Fern. Prieto, C. Aguiar, E. Días & M.I. Gut | 45_S390 | 211 | 90.17 | 113730 | 62.85 |
| *Festuca gautieri* (Hack.) K.Richt. | 46_S391 | 216 | 92.31 | 134823 | 74.51 |
| *Festuca gigantea* (L.) Vill. | 47_S392 | 211 | 90.17 | 127779 | 70.62 |
| *Festuca glauca* Vill. | 48_S393 | 217 | 92.74 | 130368 | 72.05 |
| *Festuca glumosa* Hack. ex E.B.Alexeev | 49_S394 | 215 | 91.88 | 132012 | 72.96 |
| *Festuca gracilior* (Hack.) Markgr.-Dann | 50_S396 | 221 | 94.44 | 148236 | 81.92 |
| *Festuca gracillima* Hook. F. | 51_S397 | 212 | 90.60 | 117126 | 64.73 |
| *Festuca gudoschnikovii* Stepanov | 52_S398 | 211 | 90.17 | 117858 | 65.13 |
| *Festuca henriquesii* Hack. | 53_S399 | 217 | 92.74 | 137850 | 76.18 |
| *Festuca hephaestophilla* (Nees) Nees. | 54_S400 | 187 | 79.91 | 89247 | 49.32 |
| *Festuca hieronymi* Hack. | 55_S401 | 221 | 94.44 | 124509 | 68.81 |
| *Festuca holubii* Stančík | 56_S402 | 221 | 94.44 | 135822 | 75.06 |
| *Festuca humilior* Nees | 57_S403 | 217 | 92.74 | 111735 | 61.75 |
| *Festuca hystrix* Boiss. | 58_S404 | 211 | 90.17 | 119640 | 66.12 |
| *Festuca iberica* (Hack.) K.Richt. | 59_S405 | 226 | 96.58 | 141900 | 78.42 |
| *Festuca imbaburensis* Stančik | 60_S407 | 209 | 89.32 | 131592 | 72.72 |
| *Festuca indigesta* Boiss. | 61_S408 | 221 | 94.44 | 136242 | 75.29 |
| *Festuca kolesnikovii* Tzvelev | 63_S410 | 226 | 96.58 | 138006 | 76.27 |
| *Festuca kurtziana* St.-Yves | 64_S411 | 180 | 76.92 | 74595 | 41.23 |
| *Festuca laegaardii* Stančik | 65_S412 | 218 | 93.16 | 129606 | 71.63 |
| *Festuca laevigata* Gaudin | 66_S413 | 218 | 93.16 | 137346 | 75.90 |
| *Festuca lasto* Boiss*.* | 67_S414 | 202 | 86.32 | 130362 | 72.04 |
| *Festuca lemanii* Bastard | 68_S415 | 218 | 93.16 | 141651 | 78.28 |
| *Festuca leptopogon* Stapf | 69_S416 | 202 | 86.32 | 105765 | 58.45 |
| *Festuca longiauriculata* Fuente, Ortúñez & Ferrero Lom. | 70_S418 | 224 | 95.73 | 140448 | 77.62 |
| *Festuca longipes* Stafp | 71_S419 | 205 | 87.61 | 116412 | 64.34 |
| *Festuca lugens* (E.Fourn.) Hitchc. ex Hern.-Xol. | 72_S420 | 149 | 63.68 | 53604 | 29.62 |
| *Festuca mairei* St.-Yves | 73_S421 | 220 | 94.02 | 134685 | 74.43 |
| *Festuca marginata* (Hack.) K. Richt. | 74_S422 | 220 | 94.02 | 141966 | 78.46 |
| *Festuca mekiste* Clayton | 75_S423 | 204 | 87.18 | 94173 | 52.04 |
| *Festuca modesta* Steud. | 76_S424 | 157 | 67.09 | 54540 | 30.14 |
| *Festuca mollissima* V.I. Krecz. & Bobrov | 77_S425 | 206 | 88.03 | 105150 | 58.11 |
| *Festuca molokaiensis* Soreng, P.M. Peterson & Catalán | 78_S426 | 220 | 94.02 | 131403 | 72.62 |
| *Festuca monguensis* Stančik | 79_S427 | 217 | 92.74 | 119895 | 66.26 |
| *Festuca muelleri* Vickery | 80_S429 | 220 | 94.02 | 120384 | 66.53 |
| *Festuca nevadensis* (Hack.) K. Richt. | 81_S430 | 225 | 96.15 | 126117 | 69.70 |
| *Festuca nigrescens* Lam. | 82_S431 | 214 | 91.45 | 131865 | 72.88 |
| *Festuca olgae* (Regel) Krivot. | 83_S432 | 194 | 82.91 | 86589 | 47.85 |
| *Festuca orthophylla* Pilg. | 84_S433 | 213 | 91.03 | 116784 | 64.54 |
| *Festuca ovina* L. | 85_S434 | 227 | 97.01 | 147270 | 81.39 |
| *Festuca pampeana* Speg. | 86_S435 | 223 | 95.30 | 135948 | 75.13 |
| *Festuca paniculata* (L.) Schinz & Thell | 87_S436 | 225 | 96.15 | 146322 | 80.87 |
| *Festuca parciflora* Swallen | 88_S437 | 207 | 88.46 | 130743 | 72.26 |
| *Festuca parvigluma* Steud. | 89_S438 | 231 | 98.72 | 131457 | 72.65 |
| *Festuca pilgeri* St.-Yves | 90_S440 | 225 | 96.15 | 140748 | 77.78 |
| *Festuca plebeia* R.Br. | 91_S441 | 221 | 94.44 | 138069 | 76.30 |
| *Festuca plicata* Hack. | 92_S442 | 218 | 93.16 | 134538 | 74.35 |
| *Festuca pratensis* Huds. | 93_S443 | 213 | 91.03 | 135522 | 74.90 |
| *Festuca procera* Kunth | 94_S444 | 218 | 93.16 | 135033 | 74.63 |
| *Festuca pseudeskia* Boiss. | 95_S445 | 219 | 93.59 | 144522 | 79.87 |
| *Festuca pumila* Wilk. | 96_S446 | 225 | 96.15 | 146733 | 81.09 |
| *Festuca pyrenaica* Reut. | 97_S447 | 208 | 88.89 | 123852 | 68.45 |
| *Festuca pyrogea* Speg. | 98_S448 | 222 | 94.87 | 131589 | 72.72 |
| *Festuca quadridentata* Kunth | 99_S449 | 162 | 69.23 | 66093 | 36.53 |
| *Festuca queriana* Litard. | 100_S3 | 193 | 82.48 | 94470 | 52.21 |
| *Festuca raddei* Enustsch. & Prob*. [sp. nova (F. aggr. rubra)]* | 101_S4 | 218 | 93.16 | 125178 | 69.18 |
| *Festuca reverchonii* Hack. | 102_S5 | 210 | 89.74 | 120141 | 66.40 |
| *Festuca richardsonii* Hook. | 103_S6 | 221 | 94.44 | 118959 | 65.74 |
| *Festuca rigidifolia* Tovar | 104_S7 | 216 | 92.31 | 110733 | 61.20 |
| *Festuca rubra* L. | 105_S8 | 214 | 91.45 | 127611 | 70.52 |
| *Festuca samensis* Joch.Müll. | 106_S9 | 204 | 87.18 | 94584 | 52.27 |
| *Festuca scabra* Vahl | 107_S10 | 207 | 88.46 | 100290 | 55.43 |
| *Festuca scariosa* Pau | 108_S11 | 203 | 86.75 | 101487 | 56.09 |
| *Festuca setifolia* Steud. ex Griseb. | 109_S12 | 222 | 94.87 | 125706 | 69.47 |
| *Festuca simensis* Hochst. ex A.Rich. | 111_S15 | 203 | 86.75 | 107430 | 59.37 |
| *Festuca sodiroana* Hack. ex E.B. Alexeev | 112_S16 | 219 | 93.59 | 118851 | 65.68 |
| *Festuca spectabilis* Jan. | 114_S18 | 218 | 93.16 | 125697 | 69.47 |
| *Festuca subuliflora* Scribn. | 115_S19 | 220 | 94.02 | 122325 | 67.60 |
| *Festuca subulifolia* Benth. | 116_S20 | 211 | 90.17 | 126441 | 69.88 |
| *Festuca subverticillata* (Pers.) E.B.Alexeev | 117_S21 | 220 | 94.02 | 119271 | 65.92 |
| *Festuca superba* Parodi ex Türpe | 118_S22 | 215 | 91.88 | 119451 | 66.01 |
| *Festuca triflora* J.F. Gmel. | 119_S23 | 188 | 80.34 | 97299 | 53.77 |
| *Festuca versuta* Beal | 122_S27 | 206 | 88.03 | 107310 | 59.30 |
| *Festuca viviparoidea* Krajina ex Pavlick | 123_S28 | 224 | 95.73 | 146574 | 81.00 |
| *Festuca weberbaueri* Pilg. | 124_S29 | 220 | 94.02 | 121863 | 67.35 |
| *Festuca woronowii* Hack. | 125_S30 | 221 | 94.44 | 134157 | 74.14 |
| *Festuca yalaensis* Joch.Müll. & Catalán | 126_S31 | 226 | 96.58 | 136419 | 75.39 |
| *Festuca yvesii* Sennen & Pau | 127_S32 | 222 | 94.87 | 148761 | 82.21 |
| *Hellerochloa livida* (Kunt) Willd. ex Spreng. | 128_S33 | 226 | 96.58 | 120300 | 66.48 |
| *Lolium canariense* Steud | 129_S34 | 216 | 92.31 | 149355 | 82.54 |
| *Lolium perenne* L. | 130_S36 | 216 | 92.31 | 144774 | 80.01 |
| *Lolium rigidum* Gaudin | 132_S38 | 226 | 96.58 | 146238 | 80.82 |
| *Lolium saxatile* H. Scholz & S. Scholz | 133_S39 | 223 | 95.30 | 155034 | 85.68 |
| *Lolium temulentum* L. | 134_S40 | 224 | 95.73 | 138930 | 76.78 |
| *Megalachne berteroniana* Steud. | 135_S41 | 225 | 96.15 | 132714 | 73.34 |
| *Micropyropsis tuberosa* Romero-Zarco & Cabezudo | 136_S42 | 223 | 95.30 | 143619 | 79.37 |
| *Psilurus incurvus* (Gouan) Schinz & Thell. | 137_S43 | 227 | 97.01 | 141804 | 78.37 |
| *Vulpia muralis* (Kunth) Nees | 138_S44 | 227 | 97.01 | 151380 | 83.66 |
| *Vulpia ciliata* Dumort. | 140_S47 | 217 | 92.74 | 122163 | 67.51 |
| *Vulpia membranacea* (L.) Dumort. | 142_S49 | 210 | 89.74 | 130179 | 71.94 |
| *Vulpia sicula* (C. Presl) Link | 143_S50 | 225 | 96.15 | 148782 | 82.22 |
| *Wangenheimia lima* (L.) Trin. | 144_S51 | 225 | 96.15 | 140133 | 77.44 |
|  | 132 | 234 | 100.00 | 180946 | 100.00 |

**Supplementary Table S3**. Summary statistics from HybPhaser analysis, based on target gene capture in 132 Loliinae taxa, and diversity parameters. Taxon, Sample code (Arbor Biosciences), Ploidy level, Phylogenetic group, Number of saved loci per taxon, Percentage of saved genes per taxon (from a total of 345), Sequence length of saved genes (bp) per taxon, Allele divergence (AD), Locus heterozygosity (LH), Hybridogenic group according to AD and LH classification (see text, Figures 2b, 2c, and Supplementary Fig. S5). Asterisks indicate hybridogenic group classification different from ploidy level.

| **Sample Code** | **Taxon** | **Ploidy** | **Phylogenetic group** | **no. loci** | **% genes/taxon** | **lengtht (bp)** | **AD** | **LH** | **Hybridogenic Group** |
| --- | --- | --- | --- | --- | --- | --- | --- | --- | --- |
| 1_S1 | *F. abyssinica* | 4x | Afroalpine | 292 | 62.8 | 178725 | 3.295 | 91.43 | High Polyploid* |
| 22_S135 | *F. caprina* | 4x | Afroalpine | 321 | 70.3 | 200067 | 5.731 | 96.02 | High Polyploid* |
| 25_S168 | *F. chodatiana* | 4x | Afroalpine | 304 | 65.5 | 186327 | 3.397 | 91.02 | High Polyploid* |
| 90_S440 | *F. pilgeri* | 4x | Afroalpine | 307 | 72.6 | 206469 | 3.62 | 95.17 | High Polyploid* |
| 19_S101 | *F. californica* | 8x | American - Neozeylandic | 315 | 56.5 | 160818 | 4.159 | 95.61 | High Polyploid |
| 51_S397 | *F. gracillima* | 6x | American - Neozeylandic | 301 | 62.1 | 176547 | 3.546 | 89.81 | High Polyploid |
| 64_S411 | *F. kurtziana* | 6x | American - Neozeylandic | 259 | 42.3 | 120273 | 3.392 | 90.07 | High Polyploid |
| 106_S9 | *F. samensis* | ? | American - Vulpia - Pampas | 283 | 50.2 | 142812 | 3.403 | 93.29 | High Polyploid |
| 86_S435 | *F. pampeana* | 8x | American - Vulpia - Pampas | 316 | 72 | 204819 | 4.022 | 94.53 | High Polyploid |
| 24_S157 | *F. chimborazensis* | 6x | American I | 304 | 67.6 | 192276 | 3.829 | 91.11 | High Polyploid |
| 56_S402 | *F. holubii* | ? | American I | 318 | 71.2 | 202413 | 3.886 | 92.07 | High Polyploid |
| 60_S407 | *F. imbaburensis* | 4x | American I | 305 | 69.1 | 196476 | 3.792 | 91.69 | High Polyploid* |
| 9_S439 | *F. andicola* | 4x | American II | 311 | 67.3 | 191409 | 4.399 | 92.48 | High Polyploid* |
| 14_S46 | *F. asplundii* | 6x | American II | 317 | 69 | 196305 | 4.336 | 95.73 | High Polyploid |
| 21_S124 | *F. carazana* | ? | American II | 318 | 63.7 | 181137 | 4.308 | 96.06 | High Polyploid |
| 27_S190 | *F. compresifolia* | ? | American II | 299 | 54.4 | 154845 | 3.761 | 89.94 | High Polyploid |
| 29_S212 | *F. dasyantha* | ? | American II | 295 | 53 | 150855 | 4.176 | 95.16 | High Polyploid |
| 31_S235 | *F. distichovaginata* | ? | American II | 298 | 56.8 | 161484 | 3.979 | 96.45 | High Polyploid |
| 32_S246 | *F. dolichophylla* | ? | American II | 319 | 67.3 | 191430 | 4.504 | 94.91 | High Polyploid |
| 43_S368 | *F. flacca* | 4x | American II | 287 | 50.5 | 143637 | 3.663 | 88.51 | High Polyploid* |
| 49_S394 | *F. glumosa* | 4x | American II | 311 | 70.4 | 200253 | 4.569 | 94.04 | High Polyploid* |
| 55_S401 | *F. hieronymi* | 6x | American II | 319 | 64.8 | 184224 | 4.333 | 93.87 | High Polyploid |
| 57_S403 | *F. humilior* | ? | American II | 308 | 59.2 | 168312 | 4.473 | 96.89 | High Polyploid |
| 65_S412 | *F. laegaardii* | 4x | American II | 321 | 68.3 | 194172 | 4.345 | 92.94 | High Polyploid* |
| 79_S427 | *F. monguensis* | ? | American II | 310 | 63.3 | 179895 | 4.49 | 95.05 | High Polyploid |
| 84_S433 | *F. orthophylla* | 8x | American II | 313 | 62.4 | 177528 | 4.161 | 88.71 | High Polyploid |
| 88_S437 | *F. parciflora* | 4x | American II | 301 | 69.9 | 198666 | 4.531 | 93.89 | High Polyploid* |
| 94_S444 | *F. procera* | 4x | American II | 314 | 71.8 | 204189 | 4.706 | 93.54 | High Polyploid* |
| 104_S7 | *F. rigidifolia* | ? | American II | 304 | 58.1 | 165261 | 4.092 | 95.27 | High Polyploid |
| 109_S12 | *F. setifolia* | ? | American II | 315 | 65.8 | 187149 | 4.419 | 97.26 | High Polyploid |
| 112_S16 | *F. sodiroana* | 4x | American II | 316 | 62.2 | 176994 | 4.37 | 96.28 | High Polyploid* |
| 116_S20 | *F. subulifolia* | 4x | American II | 309 | 67.7 | 192459 | 4.227 | 91.82 | High Polyploid* |
| 122_S27 | *F. versuta* | ? | American II | 296 | 56.6 | 161112 | 3.977 | 96.37 | High Polyploid |
| 124_S29 | *F. weberbaueri* | ? | American II | 318 | 64.6 | 183852 | 4.346 | 94.19 | High Polyploid |
| 126_S31 | *F. yalaensis* | ? | American II | 323 | 72.3 | 205506 | 4.808 | 97.9 | High Polyploid |
| 128_S33 | *H. livida* | ? | American II | 320 | 63 | 179259 | 4.125 | 96.05 | High Polyploid |
| 72_S420 | *F. lugens* | 4x | Asperifolia - MCSA I | 209 | 27.8 | 79029 | 2.394 | 88.43 | Old Polyploid |
| 45_S390 | *F. francoi* | 2x | Aulaxyper | 282 | 60.1 | 171072 | 1.253 | 70.42 | Diploid |
| 53_S399 | *F. henriquesii* | 2x | Aulaxyper | 307 | 72.1 | 205086 | 1.279 | 67.91 | Diploid |
| 138_S44 | *V. muralis* | 2x | Aulaxyper | 297 | 77.7 | 220929 | 0.978 | 72.67 | Diploid |
| 100_S3 | *F. querana* | 4x | Aulaxyper | 278 | 51.7 | 146925 | 2.644 | 83.33 | Old Polyploid |
| 101_S4 | *F. raddei* | ? | Aulaxyper | 304 | 66.1 | 187863 | 3.942 | 94.36 | High Polyploid |
| 103_S6 | *F. richardsonii* | 6x | Aulaxyper | 312 | 62.1 | 176754 | 4.131 | 96.92 | High Polyploid |
| 105_S8 | *F. rubra* | 6x | Aulaxyper | 310 | 67.8 | 192912 | 3.911 | 91.85 | High Polyploid |
| 59_S405 | *F. iberica* | 6x | Aulaxyper | 320 | 72.7 | 206883 | 3.725 | 94.91 | High Polyploid |
| 81_S430 | *F. nevadensis* | 10x | Aulaxyper | 314 | 65.7 | 186816 | 3.974 | 93.87 | High Polyploid |
| 82_S431 | *F. nigrescens* | 6x | Aulaxyper | 312 | 70.6 | 200769 | 4.181 | 93.73 | High Polyploid |
| 13_S35 | *F. asperula* | ? | Australia - Tasmania | 313 | 69.6 | 198051 | 4.59 | 98.17 | High Polyploid |
| 91_S441 | *F. plebeia* | ? | Australia - Tasmania | 312 | 72.1 | 204951 | 4.527 | 98.16 | High Polyploid |
| 10_S2 | *F. argentina* | 4x | Coironhuecu | 317 | 71.7 | 203904 | 2.901 | 89.81 | High Polyploid* |
| 76_S424 | *F. modesta* | 2x | Drymanthele - Muticae | 225 | 29.9 | 84903 | 3.006 | 89.04 | High Polyploid* |
| 34_S268 | *F. drymeja* | 2x | Drymanthele- Phaeochloa | 295 | 62.6 | 178071 | 1.339 | 73.1 | Diploid |
| 67_S414 | *F. lasto* | 2x | Drymanthele - Phaeochloa | 285 | 69.4 | 197256 | 1.512 | 71.95 | Old Polyploid* |
| 118_S22 | *F. superba* | 8x | Drymanthele s. l. - MCSA I | 311 | 64.5 | 183396 | 4.651 | 92.72 | High Polyploid |
| 30_S224 | *F. dichoclada* | ? | Erosiflorae - MCSA I | 322 | 67.2 | 191049 | 4.976 | 94.85 | High Polyploid |
| 99_S449 | *F. quadridentata* | ? | Erosiflorae - MCSA II | 243 | 35.5 | 100935 | 3.285 | 89.24 | High Polyploid |
| 2_S112 | *F. acuminata* | 2x | Eskia | 310 | 76.8 | 218532 | 1.105 | 55.38 | Diploid |
| 46_S391 | *F. gautieri* | 2x | Eskia | 309 | 71.5 | 203436 | 1.156 | 66.15 | Diploid |
| 96_S446 | *F. pumila* | 2x | Eskia | 312 | 75.4 | 214479 | 1.249 | 68.28 | Diploid |
| 125_S30 | *F. woronowii* | 2x | Eskia | 306 | 69.1 | 196596 | 1.176 | 72.62 | Diploid |
| 36_S290 | *F. elegans* | 4x | Eskia | 286 | 59.1 | 167967 | 1.519 | 74.76 | Old Polyploid |
| 20_S113 | *F. capillifolia* | 2x | Exaratae - Loretia | 301 | 73.8 | 209961 | 1.005 | 55.11 | Diploid |
| 92_S442 | *F. plicata* | 2x | Exaratae - Loretia | 305 | 70.2 | 199536 | 1.146 | 70.81 | Diploid |
| 97_S447 | *F. pyrenaica* | 4x | Exaratae - Loretia | 291 | 65.6 | 186648 | 1.087 | 64.54 | Diploid* |
| 142_S49 | *V. membranacea* | 2x | Exaratae - Loretia | 309 | 69.1 | 196485 | 0.678 | 30.89 | Diploid |
| 143_S50 | *V. sicula* | 2x | Exaratae - Loretia | 303 | 76.9 | 218718 | 1.149 | 70.09 | Diploid |
| 54_S400 | *F. hephaestophila* | 4x | Exaratae - Loretia | 270 | 47.1 | 133845 | 2.249 | 79.51 | Old Polyploid |
| 117_S21 | *F. subverticillata* | 4x | Exaratae-Loretia | 314 | 63.4 | 180321 | 4.437 | 96.31 | High Polyploid* |
| 135_S41 | *M. berteroniana* | ? | Fernandezian | 314 | 69.7 | 198327 | 3.699 | 95.47 | High Polyploid |
| 5_S395 | *F. alpina* | 2x | Festuca | 300 | 67.3 | 191439 | 0.889 | 57.05 | Diploid |
| 41_S346 | *F. filiformis* | 2x | Festuca | 305 | 72.9 | 207282 | 1.288 | 67.78 | Diploid |
| 58_S404 | *F. hystrix* | 2x | Festuca | 301 | 63.6 | 181002 | 1.283 | 72.47 | Diploid |
| 63_S410 | *F. kolesnikovii* | ? | Festuca | 314 | 71.3 | 202914 | 1.078 | 68.39 | Diploid |
| 70_S418 | *F. longiauriculata* | 2x | Festuca | 317 | 72.6 | 206487 | 1.446 | 77.71 | Diploid |
| 74_S422 | *F. marginata* | 2x | Festuca | 308 | 74.2 | 211047 | 1.291 | 69.44 | Diploid |
| 77_S425 | *F. mollissima* | 2x | Festuca | 293 | 55.7 | 158544 | 1.202 | 69.38 | Diploid |
| 85_S434 | *F. ovina* | 2x | Festuca | 316 | 75.6 | 215052 | 1.407 | 77.48 | Diploid |
| 102_S5 | *F. reverchonii* | 2x | Festuca | 283 | 62.7 | 178215 | 1.285 | 76.92 | Diploid |
| 16_S68 | *F. brevipila* | 6x | Festuca | 253 | 48.1 | 136776 | 1.986 | 80.74 | Old Polyploid* |
| 48_S393 | *F. glauca* | 2x | Festuca | 313 | 69.2 | 196851 | 2.314 | 87.35 | Old Polyploid* |
| 50_S396 | *F. gracilior* | 4x | Festuca | 313 | 76.7 | 218181 | 1.987 | 86.54 | Old Polyploid |
| 144_S51 | *W. lima* | 2x | Festuca | 308 | 73.2 | 208191 | 1.715 | 87.31 | Old Polyploid* |
| 61_S408 | *F. indigesta* | 6x | Festuca | 312 | 70.8 | 201399 | 2.53 | 92.94 | High Polyploid |
| 66_S413 | *F. laevigata* | 8x | Festuca | 310 | 72.5 | 206205 | 2.332 | 91.93 | High Polyploid |
| 68_S415 | *F. lemanii* | 6x | Festuca | 309 | 74.4 | 211680 | 2.395 | 91.95 | High Polyploid |
| 98_S448 | *F. pyrogea* | ? | Festuca | 307 | 69.1 | 196641 | 2.498 | 90.88 | High Polyploid |
| 127_S32 | *F. yvesii* | 8x | Festuca | 312 | 77 | 218916 | 2.421 | 93.01 | High Polyploid |
| 123_S28 | *F. viviparoidea* | 8x | Festuca | 313 | 75.8 | 215727 | 2.385 | 94.55 | High Polyploid |
| 18_S90 | *F. caldasii* | 4x | Glabricarpae - MCSA I | 291 | 61.1 | 173913 | 2.505 | 80.13 | Old Polyploid |
| 83_S432 | *F. olgae* | 4x | Leucopoa | 284 | 46 | 130845 | 2.472 | 89.66 | Old Polyploid |
| 6_S406 | *F. altaica* | 4x | Leucopoa | 315 | 73.7 | 209700 | 3.076 | 93.56 | High Polyploid* |
| 17_S79 | *F. calabrica* | ? | Leucopoa | 311 | 71.7 | 204012 | 3.33 | 93.81 | High Polyploid |
| 80_S429 | *F. muelleri* | ? | Leucopoa | 319 | 64.3 | 182841 | 4.107 | 96.6 | High Polyploid |
| 114_S18 | *F. spectabilis* | 6x | Leucopoa | 311 | 66.9 | 190230 | 3.15 | 94.72 | High Polyploid |
| 26_S179 | *F. coerulescens* | 2x | Lojaconoa | 294 | 75.2 | 214023 | 1.235 | 77.61 | Diploid |
| 119_S23 | *F. triflora* | 2x | Lojaconoa | 282 | 53.5 | 152232 | 0.863 | 39.37 | Diploid |
| 95_S445 | *F. pseudeskia* | 2x | Pseudoscariosa | 308 | 75.3 | 214113 | 0.956 | 55.52 | Diploid |
| 137_S43 | *P. incurvus* | 4x | Psilurus - Vulpia | 314 | 74 | 210390 | 3.415 | 94.31 | High Polyploid* |
| 140_S47 | *V. ciliata* | 4x | Psilurus - Vulpia | 309 | 65.2 | 185526 | 3.075 | 92.52 | High Polyploid* |
| 8_S428 | *F. amplissima* | 6x | Ruprechtia - MCSA II | 309 | 64.7 | 184068 | 2.768 | 87.77 | Old Polyploid* |
| 23_S146 | *F. valdesii* | ? | Ruprechtia - MCSA II | 317 | 62.9 | 178797 | 3.765 | 94.74 | High Polyploid |
| 108_S11 | *F. scariosa* | 2x | Scariosae | 284 | 54.3 | 154386 | 1.438 | 68.33 | Diploid |
| 44_S379 | *F. fontqueri* | 2x | Schedonorus - Lolium | 309 | 75.9 | 215892 | 0.934 | 58.59 | Diploid |
| 93_S443 | *F. pratensis* | 2x | Schedonorus - Lolium | 302 | 72 | 204855 | 0.935 | 59.87 | Diploid |
| 129_S34 | *L. canariense* | 2x | Schedonorus - Lolium | 318 | 77.4 | 220212 | 0.504 | 22.36 | Diploid |
| 133_S39 | *L. saxatile* | 2x | Schedonorus - Lolium | 319 | 79.7 | 226599 | 0.516 | 26.36 | Diploid |
| 134_S40 | *L. temulentum* | 2x | Schedonorus - Lolium | 316 | 72.3 | 205494 | 0.527 | 38.97 | Diploid |
| 136_S42 | *M. tuberosa* | 2x | Schedonorus - Lolium | 300 | 75.2 | 213918 | 0.815 | 55.45 | Diploid |
| 111_S15 | *F. simensis* | 4x | Schedonorus - Lolium | 290 | 57.9 | 164628 | 1.937 | 84.59 | Old Polyploid |
| 130_S36 | *L. perenne* | 2x | Schedonorus - Lolium | 300 | 75.5 | 214821 | 1.776 | 83.44 | Old Polyploid* |
| 11_S13 | *F. atlantigena* | 8x | Schedonorus - Lolium | 306 | 69.2 | 196863 | 3.064 | 93.75 | High Polyploid |
| 33_S257 | *F. dracomontana* | ? | Schedonorus - Lolium | 309 | 55.4 | 157683 | 3.246 | 92.72 | High Polyploid |
| 40_S335 | *F. fenas* | 4x | Schedonorus - Lolium | 302 | 65.2 | 185553 | 2.663 | 92.77 | High Polyploid* |
| 47_S392 | *F. gigantea* | 6x | Schedonorus - Lolium | 300 | 68.1 | 193779 | 2.596 | 93.97 | High Polyploid |
| 52_S398 | *F. gudoschnikovii* | 4x | Schedonorus - Lolium | 305 | 62.7 | 178263 | 2.513 | 92.09 | High Polyploid* |
| 12_S24 | *F. letourneuxiana* | 10x | Schedonorus - Lolium | 307 | 68 | 193500 | 2.944 | 93.17 | High Polyploid |
| 73_S421 | *F. mairei* | 4x | Schedonorus - Lolium | 314 | 70.5 | 200493 | 2.698 | 93.25 | High Polyploid* |
| 132_S38 | *L. rigidum* | 2x | Schedonorus - Lolium | 319 | 75.7 | 215232 | 2.345 | 92.49 | High Polyploid* |
| 35_S279 | *F. durandoi* | 4x | Subbulbosae | 297 | 65.3 | 185805 | 1.2 | 59.09 | Diploid |
| 87_S436 | *F. paniculata* | 2x | Subbulbosae | 306 | 75.4 | 214416 | 1.174 | 71.21 | Diploid |
| 4_S334 | *F. aloha* | ? | Subulatae-Hawaiian | 307 | 54 | 153726 | 3.426 | 92.06 | High Polyploid |
| 39_S323 | *F. extremiorientalis* | 4x | Subulatae - Hawaiian | 319 | 64.2 | 182601 | 3.53 | 92.07 | High Polyploid* |
| 69_S416 | *F. leptopogon* | 4x | Subulatae - Hawaiian | 290 | 57.2 | 162759 | 3.197 | 85.38 | High Polyploid* |
| 78_S426 | *F. molokaiensis* | ? | Subulatae - Hawaiian | 319 | 69.1 | 196449 | 3.733 | 88.45 | High Polyploid |
| 89_S438 | *F. parvigluma* | 4x | Subulatae - Hawaiian | 327 | 68.2 | 193911 | 3.651 | 92.28 | High Polyploid* |
| 115_S19 | *F. subuliflora* | 4x | Subuliflorae | 318 | 65.1 | 185079 | 3.247 | 89.2 | High Polyploid* |
| 3_S223 | *F. africana* | 10x | Tropical-South African | 250 | 43.1 | 122445 | 2.446 | 76.65 | Old Polyploid* |
| 107_S10 | *F. scabra* | 4x | Tropical - South African | 295 | 54.4 | 154797 | 2.72 | 81.79 | Old Polyploid |
| 28_S201 | *F. costata* | 4x | Tropical - South African | 310 | 73 | 207591 | 3.729 | 95.43 | High Polyploid* |
| 37_S301 | *F. engleri* | ? | Tropical - South African | 306 | 64.5 | 183534 | 3.137 | 92.52 | High Polyploid |
| 71_S419 | *F. longipes* | ? | Tropical - South African | 302 | 63.2 | 179652 | 3.661 | 86.5 | High Polyploid |
| 75_S423 | *F. mekiste* | ? | Tropical - South African | 296 | 50.7 | 144108 | 4.372 | 94.08 | High Polyploid |

**Supplementary Table S4.** Values of site-specific Concordance Factor (sCF) and gene-specific Concordance Factor (gCF) calculated in IQtree2 on node branches of the ML single-copy gene phylogeny of Loliinae constructed from the concatenated supermatrix of the scg-strict data set (see Figure 1), where sCF corresponds to the concordance at individual sites in sequences and gCF to the concordance at the gene level.

| **Node Number** | **Site-specific Concordance Factor (sCF)** | **Gene-specific Concordance Factor (gCF)** |
| --- | --- | --- |
| 136 | 43.92 | 3.82 |
| 137 | 36.27 | 1.76 |
| 138 | 46.32 | 1.65 |
| 139 | 34.28 | 1.09 |
| 140 | 47.72 | 1.49 |
| 141 | 56.48 | 1.32 |
| 142 | 34.04 | 0.87 |
| 143 | 29.42 | 0.44 |
| 144 | 31.82 | 3.09 |
| 145 | 37.9 | 21.03 |
| 146 | 64.89 | 2.42 |
| 147 | 41.43 | 4.28 |
| 148 | 52.17 | 16.35 |
| 149 | 47.92 | 18.33 |
| 150 | 30.62 | 6.64 |
| 151 | 18.18 | 8 |
| 152 | 17.2 | 5.62 |
| 153 | 26.22 | 3.35 |
| 154 | 31.06 | 3.41 |
| 155 | 40.25 | 14.08 |
| 156 | 33.26 | 16.92 |
| 157 | 48.67 | 17.65 |
| 158 | 70.83 | 13.2 |
| 159 | 31.16 | 24.4 |
| 160 | 57.68 | 2.28 |
| 161 | 33.94 | 4.52 |
| 162 | 25.94 | 38.55 |
| 163 | 59.75 | 15.98 |
| 164 | 73.14 | 0 |
| 165 | 71.99 | 4.17 |
| 166 | 22.71 | 2.23 |
| 167 | 32.33 | 10.18 |
| 168 | 33.42 | 20.83 |
| 169 | 14.96 | 16.22 |
| 170 | 47.43 | 0.85 |
| 171 | 87.45 | 73.52 |
| 172 | 47.41 | 57.69 |
| 173 | 42.98 | 1.29 |
| 174 | 52 | 0.42 |
| 175 | 37.12 | 0 |
| 176 | 33.9 | 0 |
| 177 | 32.96 | 0.9 |
| 178 | 27.04 | 0 |
| 179 | 33.15 | 0 |
| 180 | 33.21 | 0.44 |
| 181 | 25.2 | 2.4 |
| 182 | 38.29 | 4.25 |
| 183 | 31.28 | 0.43 |
| 184 | 37.87 | 0.88 |
| 185 | 38.13 | 1.83 |
| 186 | 31.21 | 0.46 |
| 187 | 45.89 | 4.55 |
| 188 | 51.9 | 23.15 |
| 189 | 61.62 | 7.14 |
| 190 | 68.86 | 16.91 |
| 191 | 51.93 | 1.35 |
| 192 | 47.3 | 4.48 |
| 193 | 43.95 | 0.5 |
| 194 | 32.58 | 4.95 |
| 195 | 44.18 | 32.72 |
| 196 | 68.76 | 1.48 |
| 197 | 33.05 | 5.13 |
| 198 | 48.43 | 11.11 |
| 199 | 73.12 | 0 |
| 200 | 39.52 | 0 |
| 201 | 40.4 | 9.71 |
| 202 | 59.12 | 0.87 |
| 203 | 39.17 | 2.94 |
| 204 | 56.49 | 6.38 |
| 205 | 55.12 | 4.9 |
| 206 | 25.99 | 13.87 |
| 207 | 66.83 | 0.88 |
| 208 | 44.55 | 3.94 |
| 209 | 26.2 | 16.57 |
| 210 | 60.1 | 44.63 |
| 211 | 84.7 | 15.82 |
| 212 | 59.96 | 0 |
| 213 | 26.29 | 0 |
| 214 | 35.99 | 1.01 |
| 215 | 36.73 | 6.45 |
| 216 | 56.93 | 10.59 |
| 217 | 57.38 | 0.72 |
| 218 | 20.53 | 15.04 |
| 219 | 65.69 | 0.45 |
| 220 | 49.27 | 0 |
| 221 | 62.65 | 4.03 |
| 222 | 26.44 | 1.94 |
| 223 | 48.33 | 1.6 |
| 224 | 39.56 | 12.28 |
| 225 | 49.11 | 0 |
| 226 | 33.28 | 0 |
| 227 | 41.87 | 0 |
| 228 | 30.18 | 0 |
| 229 | 55.25 | 0 |
| 230 | 32.32 | 0 |
| 231 | 34.49 | 0.58 |
| 232 | 40.08 | 4.03 |
| 233 | 42.66 | 0 |
| 234 | 37.09 | 0.54 |
| 235 | 34.66 | 2.68 |
| 236 | 30.04 | 0 |
| 237 | 51.89 | 1.83 |
| 238 | 41.51 | 2.14 |
| 239 | 63.51 | 0 |
| 240 | 34.59 | 0 |
| 241 | 37.34 | 11.04 |
| 242 | 58.02 | 7.3 |
| 243 | 58.56 | 0.95 |
| 244 | 55.98 | 1.14 |
| 245 | 47.59 | 4.2 |
| 246 | 42.43 | 4.86 |
| 247 | 55.52 | 0 |
| 248 | 28.39 | 0 |
| 249 | 36.34 | 0 |
| 250 | 27.36 | 0 |
| 251 | 42.83 | 2.76 |
| 252 | 50.77 | 5.26 |
| 253 | 51.86 | 2.73 |
| 254 | 59.59 | 1.88 |
| 255 | 26.06 | 10.95 |
| 256 | 27.55 | 11.73 |
| 257 | 31.02 | 12.63 |
| 258 | 69.75 | 29.5 |
| 259 | 78.38 | 6.83 |
| 260 | 47.01 | 4.61 |
| 261 | 35.34 | 10.45 |
| 262 | 56.87 | 3.26 |
| 263 | 30.53 | 41.14 |
| 264 | 68.75 | 10.64 |
| 265 | 51.89 | 17.95 |
| 266 | 35.64 | 2.7 |
| 267 | 47.58 | 11.59 |

**Supplementary Table S5.** Quartet support values of nodes in the ASTRAL-III phylogeny of Loliinae based on 234 nuclear single-copy gene trees. Node number and Proportion of quartets in the gene trees that agree with the species tree for the main topology (q1), the first alternative topology (q2), and the second alternative topology (q3) per node. Minimum (min), maximum (max), and mean values of q1, q2, and q3 values across the topology.

| **Node Number** | **Q1** | **Q2** | **Q3** |
| --- | --- | --- | --- |
| 136 | NA | NA | NA |
| 137 | NA | NA | NA |
| 138 | 64.08 | 12.64 | 23.28 |
| 139 | 94.26 | 2.79 | 2.95 |
| 140 | 39.69 | 35.24 | 25.07 |
| 141 | 39.48 | 28.33 | 32.19 |
| 142 | 42.57 | 34.15 | 23.28 |
| 143 | 43.93 | 28.53 | 27.55 |
| 144 | 53.54 | 28.86 | 17.60 |
| 145 | 41.91 | 26.14 | 31.95 |
| 146 | 39.56 | 37.46 | 22.97 |
| 147 | 45.85 | 27.10 | 27.05 |
| 148 | 38.06 | 32.97 | 28.97 |
| 149 | 35.34 | 29.87 | 34.80 |
| 150 | 69.00 | 19.63 | 11.37 |
| 151 | 39.98 | 25.78 | 34.24 |
| 152 | 40.36 | 26.77 | 32.87 |
| 153 | 40.39 | 33.51 | 26.11 |
| 154 | 36.70 | 32.61 | 30.69 |
| 155 | 39.02 | 27.80 | 33.18 |
| 156 | 37.69 | 29.57 | 32.73 |
| 157 | 41.78 | 34.82 | 23.40 |
| 158 | 30.16 | 36.95 | 32.89 |
| 159 | 48.24 | 25.66 | 26.10 |
| 160 | 36.90 | 27.46 | 35.64 |
| 161 | 44.65 | 29.14 | 26.21 |
| 162 | 35.82 | 28.35 | 35.83 |
| 163 | 38.41 | 30.66 | 30.94 |
| 164 | 62.03 | 21.27 | 16.69 |
| 165 | 35.91 | 31.28 | 32.81 |
| 166 | 47.23 | 24.37 | 28.40 |
| 167 | 36.33 | 29.79 | 33.88 |
| 168 | 71.70 | 13.77 | 14.52 |
| 169 | 38.95 | 28.67 | 32.38 |
| 170 | 42.58 | 33.87 | 23.55 |
| 171 | 36.34 | 30.99 | 32.67 |
| 172 | 43.66 | 25.27 | 31.07 |
| 173 | 44.96 | 29.39 | 25.65 |
| 174 | 46.36 | 34.26 | 19.38 |
| 175 | 45.53 | 22.95 | 31.52 |
| 176 | 47.65 | 29.58 | 22.77 |
| 177 | 40.37 | 23.64 | 35.99 |
| 178 | 37.96 | 30.00 | 32.05 |
| 179 | 36.89 | 35.03 | 28.08 |
| 180 | 46.80 | 30.47 | 22.73 |
| 181 | 49.60 | 23.09 | 27.31 |
| 182 | 37.42 | 35.35 | 27.23 |
| 183 | 49.49 | 26.30 | 24.21 |
| 184 | 38.71 | 34.57 | 26.72 |
| 185 | 35.91 | 31.85 | 32.24 |
| 186 | 47.73 | 30.69 | 21.58 |
| 187 | 39.93 | 29.00 | 31.07 |
| 188 | 43.54 | 33.13 | 23.32 |
| 189 | 48.46 | 28.36 | 23.18 |
| 190 | 44.71 | 31.99 | 23.31 |
| 191 | 33.59 | 37.54 | 28.86 |
| 192 | 36.30 | 26.48 | 37.22 |
| 193 | 44.68 | 30.01 | 25.31 |
| 194 | 38.36 | 36.37 | 25.27 |
| 195 | 39.59 | 31.85 | 28.56 |
| 196 | 34.95 | 34.17 | 30.88 |
| 197 | 35.85 | 34.04 | 30.11 |
| 198 | 38.92 | 33.19 | 27.88 |
| 199 | 36.44 | 31.97 | 31.59 |
| 200 | 36.10 | 29.64 | 34.27 |
| 201 | 36.21 | 31.71 | 32.08 |
| 202 | 34.94 | 34.16 | 30.91 |
| 203 | 38.11 | 36.51 | 25.38 |
| 204 | 36.42 | 33.79 | 29.79 |
| 205 | 34.82 | 30.45 | 34.73 |
| 206 | 40.88 | 31.22 | 27.90 |
| 207 | 36.09 | 32.79 | 31.12 |
| 208 | 45.04 | 29.35 | 25.61 |
| 209 | 35.71 | 33.64 | 30.66 |
| 210 | 38.74 | 33.23 | 28.03 |
| 211 | 33.83 | 33.81 | 32.36 |
| 212 | 34.96 | 34.47 | 30.57 |
| 213 | 35.75 | 34.10 | 30.15 |
| 214 | 35.57 | 32.97 | 31.46 |
| 215 | 34.54 | 32.48 | 32.98 |
| 216 | 37.19 | 32.17 | 30.64 |
| 217 | 33.92 | 33.79 | 32.29 |
| 218 | 36.21 | 31.25 | 32.55 |
| 219 | 35.21 | 32.13 | 32.65 |
| 220 | 37.23 | 33.22 | 29.54 |
| 221 | 35.01 | 33.48 | 31.51 |
| 222 | 38.27 | 34.37 | 27.36 |
| 223 | 40.12 | 23.22 | 36.66 |
| 224 | 37.68 | 29.38 | 32.94 |
| 225 | 36.84 | 31.79 | 31.37 |
| 226 | 38.09 | 28.97 | 32.94 |
| 227 | 48.39 | 23.95 | 27.66 |
| 228 | 35.33 | 33.17 | 31.50 |
| 229 | 45.38 | 25.75 | 28.88 |
| 230 | 42.24 | 25.00 | 32.77 |
| 231 | 37.88 | 32.31 | 29.82 |
| 232 | 44.34 | 23.59 | 32.07 |
| 233 | 38.07 | 31.05 | 30.88 |
| 234 | 45.35 | 25.37 | 29.28 |
| 235 | 47.56 | 23.81 | 28.62 |
| 236 | 33.04 | 33.73 | 33.24 |
| 237 | 37.60 | 36.82 | 25.58 |
| 238 | 43.64 | 25.98 | 30.38 |
| 239 | 63.00 | 17.68 | 19.32 |
| 240 | 36.34 | 35.80 | 27.87 |
| 241 | 42.19 | 25.94 | 31.87 |
| 242 | 37.62 | 29.51 | 32.87 |
| 243 | 35.10 | 29.97 | 34.93 |
| 244 | 40.29 | 27.75 | 31.96 |
| 245 | 40.23 | 34.88 | 24.89 |
| 246 | 43.11 | 34.09 | 22.80 |
| 247 | 41.40 | 36.68 | 21.92 |
| 248 | 40.25 | 35.52 | 24.23 |
| 249 | 36.09 | 32.95 | 30.97 |
| 250 | 40.88 | 34.12 | 25.00 |
| 251 | 35.15 | 30.46 | 34.39 |
| 252 | 44.88 | 18.68 | 36.44 |
| 253 | 57.33 | 22.39 | 20.29 |
| 254 | 38.00 | 26.44 | 35.56 |
| 255 | 44.58 | 34.16 | 21.26 |
| 256 | 35.24 | 30.78 | 33.99 |
| 257 | 39.09 | 33.40 | 27.51 |
| 258 | 36.10 | 33.20 | 30.70 |
| 259 | 37.35 | 29.79 | 32.86 |
| 260 | 38.20 | 30.77 | 31.03 |
| 261 | 60.37 | 21.53 | 18.10 |
| 262 | 37.92 | 32.38 | 29.71 |
| 263 | 47.25 | 26.31 | 26.44 |
| 264 | 35.07 | 30.12 | 34.81 |
| 265 | 43.33 | 27.45 | 29.22 |
| 266 | 37.44 | 33.51 | 29.06 |
| 267 | 36.10 | 32.90 | 31.00 |
| 268 | 35.25 | 34.64 | 30.11 |
| 269 | 52.77 | 23.73 | 23.50 |
| **min** | 30.16 | 2.79 | 2.95 |
| **max** | 94.26 | 37.54 | 37.22 |
| **mean** | 41.46 | 29.87 | 28.66 |

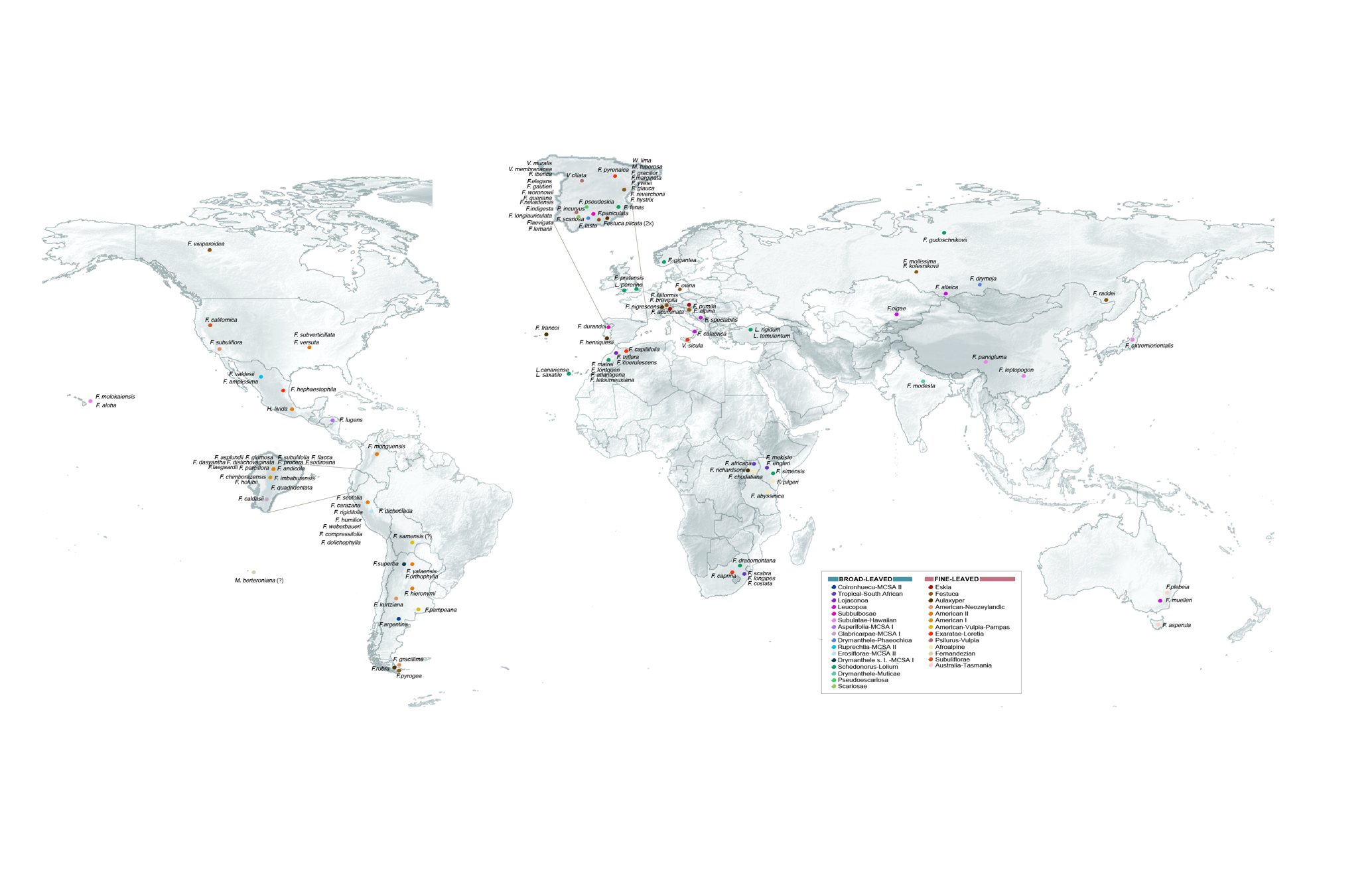

**Supplementary Figure S1**. World map showing the geographic sampling of the Loliinae taxa analyzed in this study (see also Supplementary Table S1).

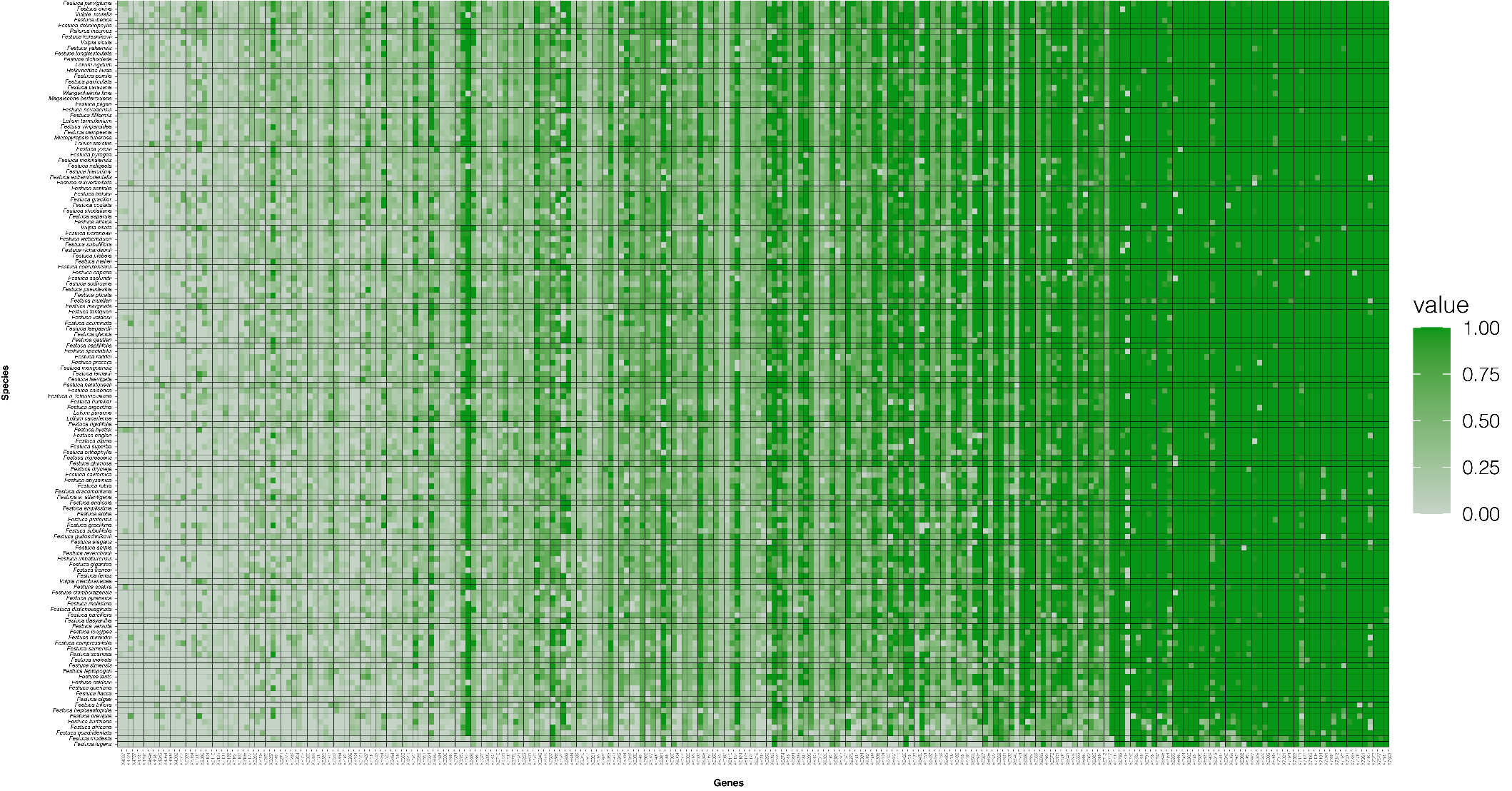

**Supplementary Figure S2**. Density statistics plot of single-copy nuclear genes processed by HybPiper in the 132 Loliinae samples studied.

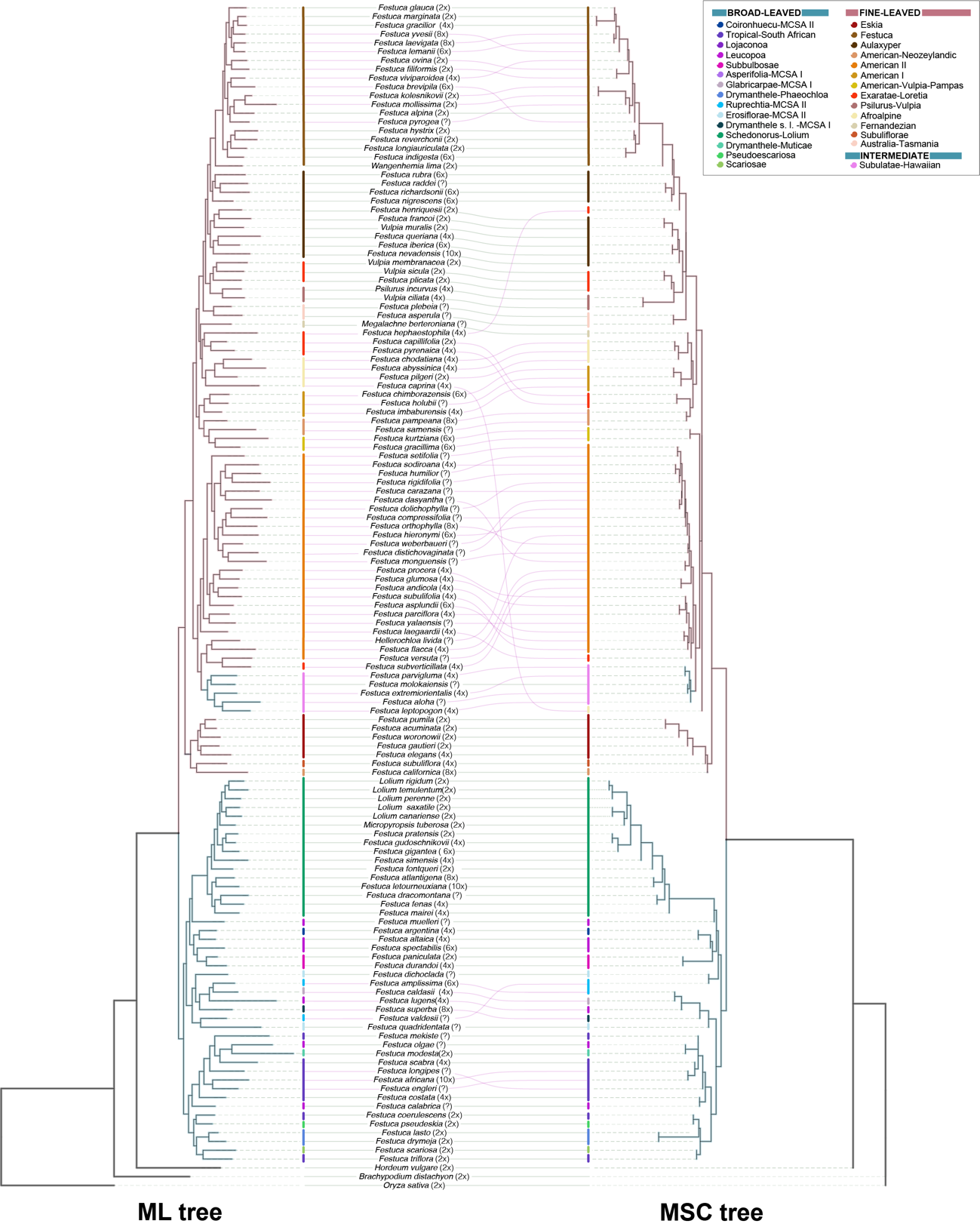
**Supplementary Figure S3**. Contrasted topologies of the nuclear Loliinae scg-strict supermatrix ML (a) and MSC (b) trees based on 234 genes and 132 taxa. The two topologies show overall congruent relationships except for the lineages and species highlighted in pink.

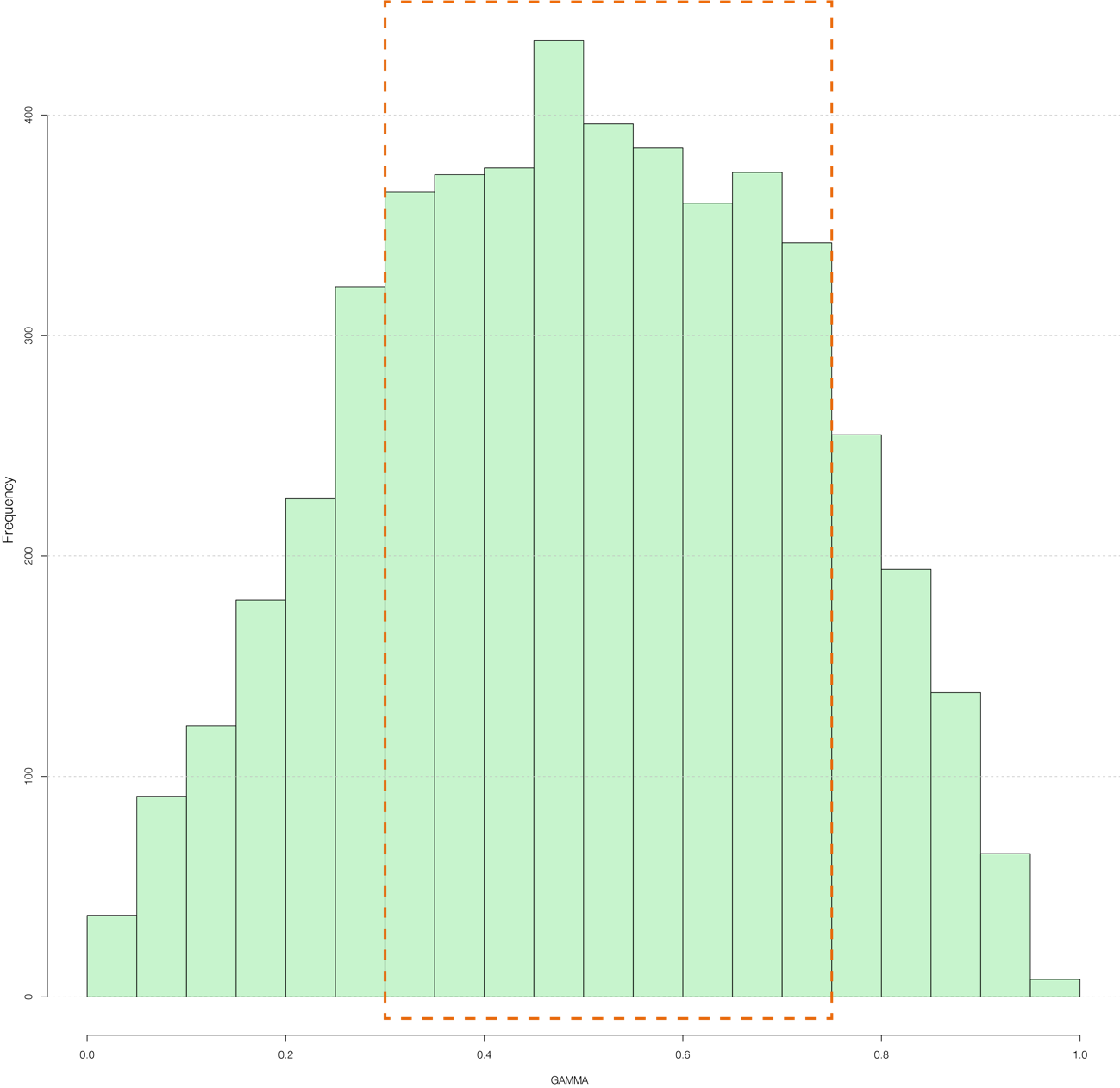

**Supplementary Figure S4**. Hybridization level among 132 studied Loliinae taxa estimated by HyDe. Histogram of frequencies of gamma values showing the highest frequencies for gamma scores of 0.3 to 0.7.

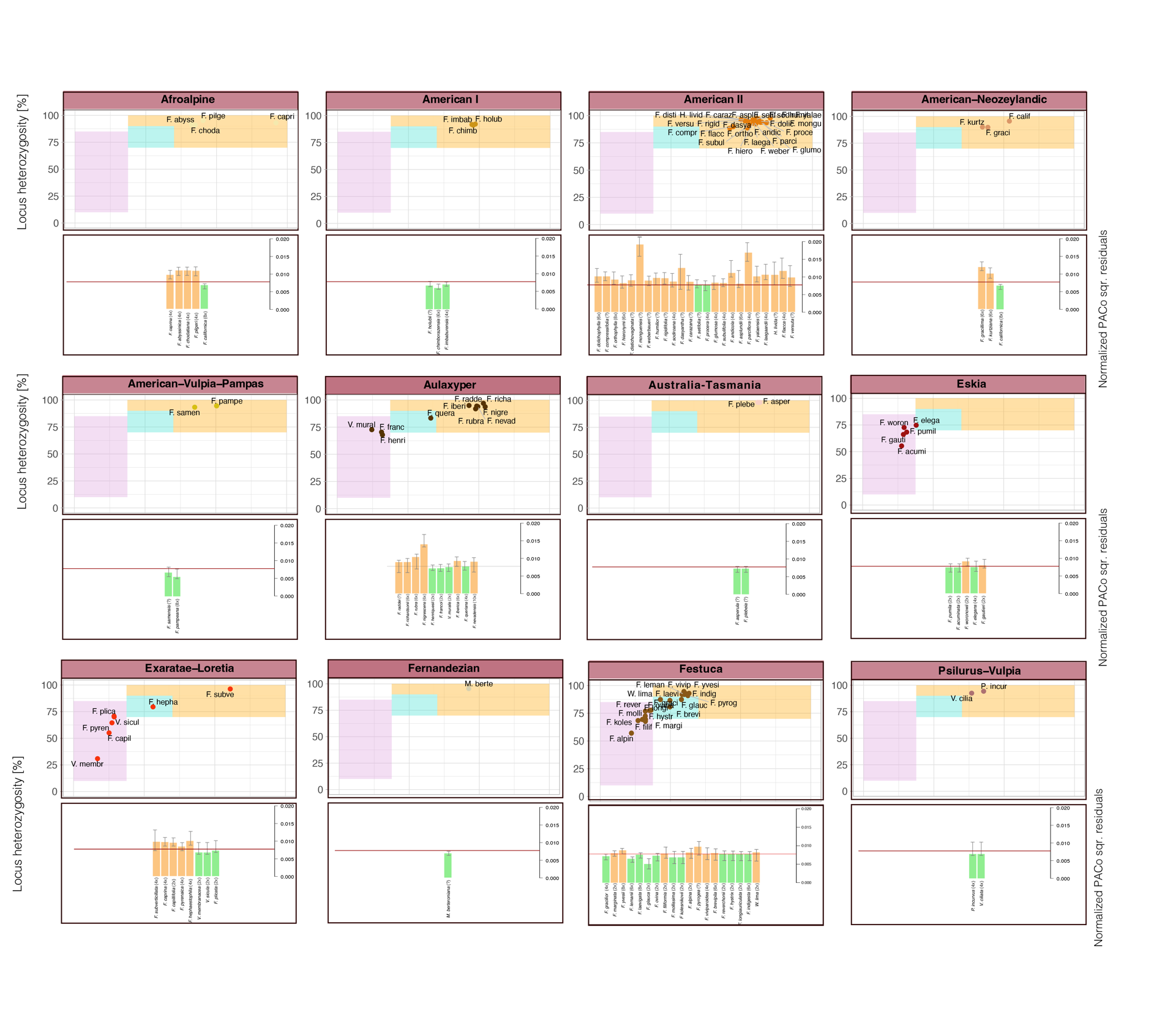

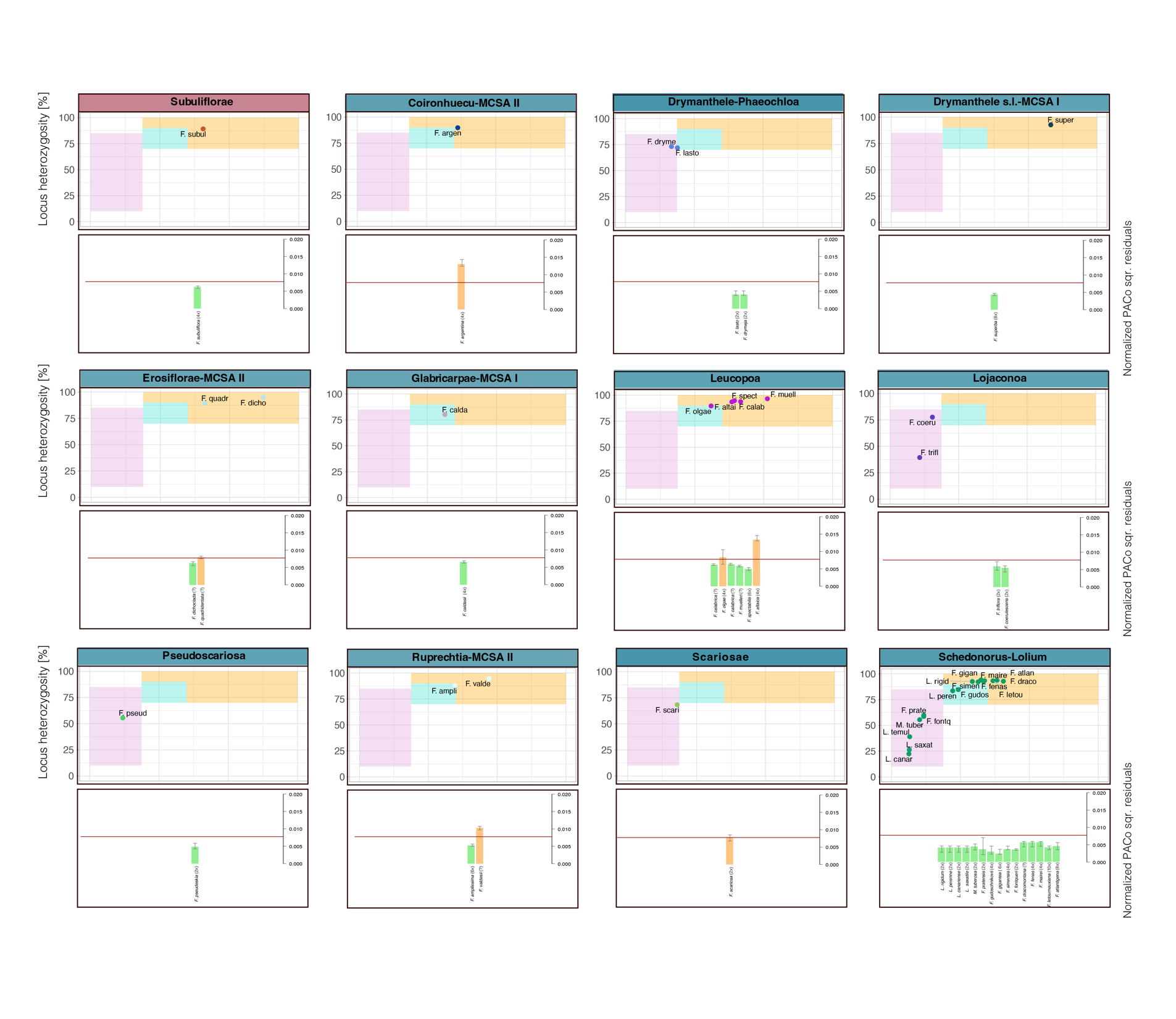

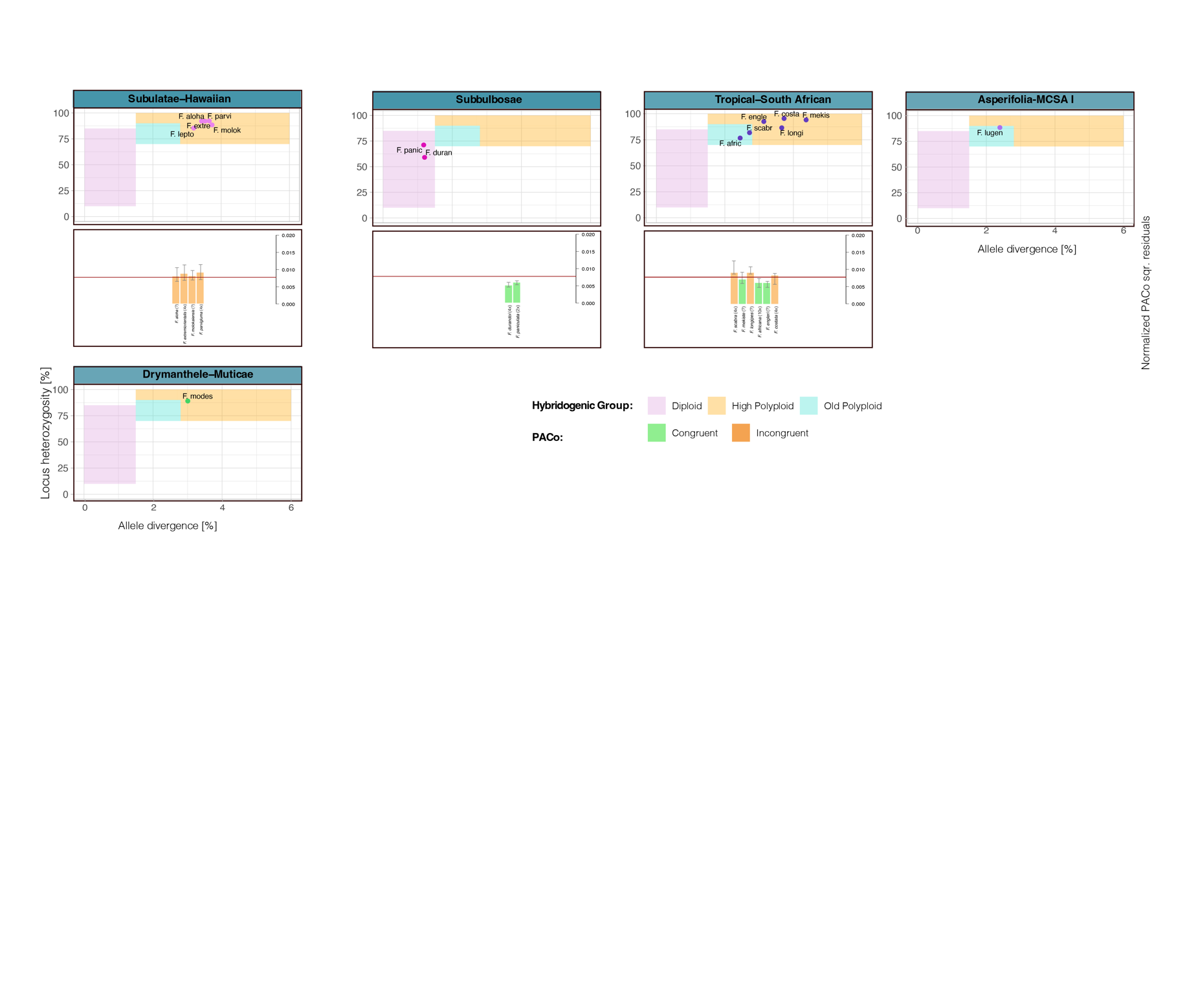

**Supplementary Figure S5**. Scatter plots of locus heterozygosity (LH) and allele divergence (AD) of the main groups of Loliinae estimated by HybPhaser (top) and box plots showing normalized squared residuals of PACo obtained from 1,000 random replicates of nuclear-plastome associations for each taxon of the group (below). LH and AD values correspond to the per-sample means of all available genes for a sample (see also Supplementary Table S3). Colored ranges indicate the three hybridogenic classes (pink: diploid, turquoise: old polyploids; yellow: highly polyploid). In the PACo boxplots, the median values of normalized squared residuals above the threshold line (0.0077) are linked to species showing incongruence between nuclear and plastome-derived trees (orange boxes) while values below the threshold correspond to topologically congruent species (green boxes) (see Figure 4b).

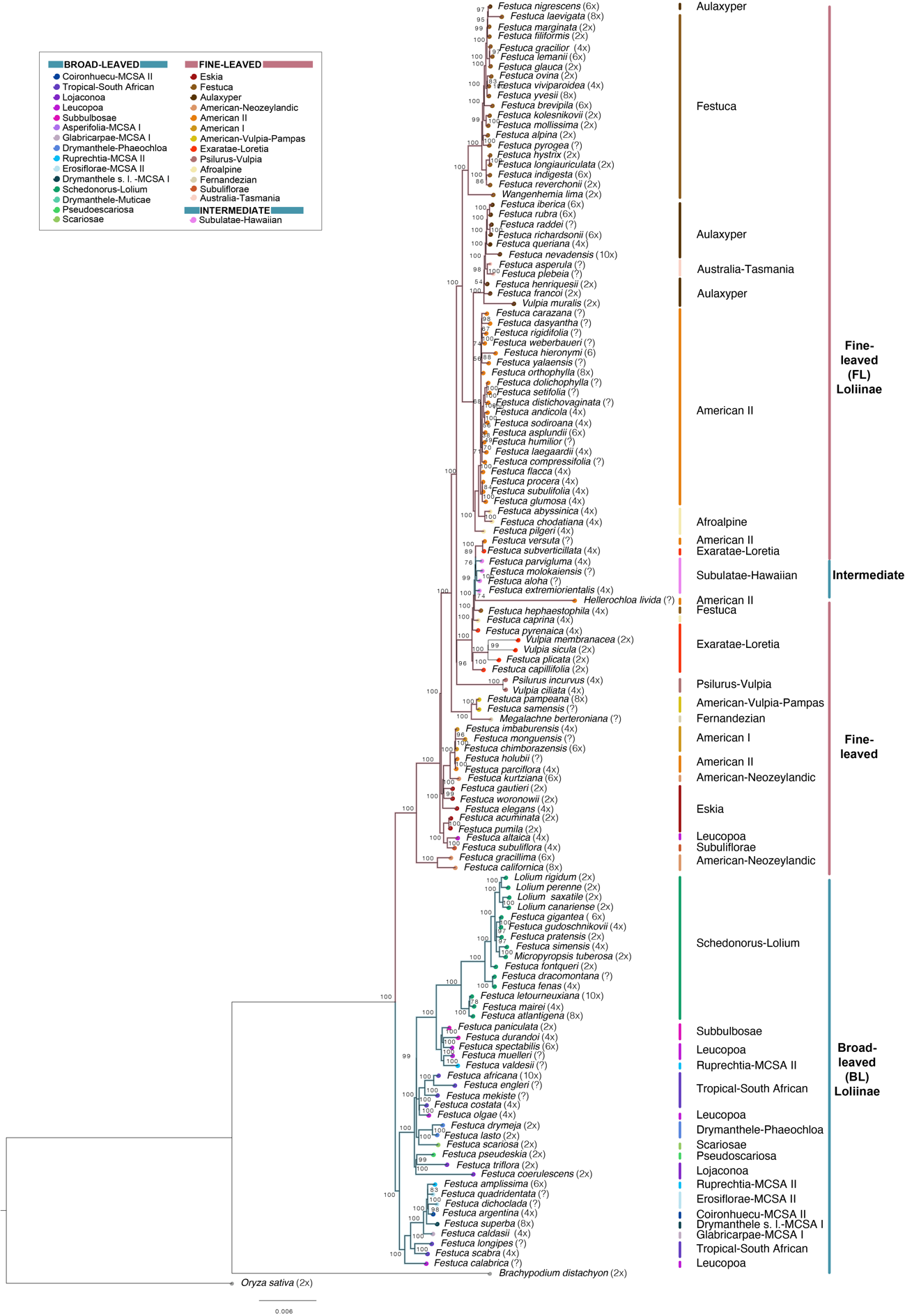

**Supplementary Figure S6.** Maximum likelihood plastome tree of Loliinae computed with Iqtree2 showing the relationships among the 128 studied taxa. Numbers on branches indicate UltraFast Bootstrap support (BS) values. *Oryza sativa* was used to root the tree. Color codes of Loliinae lineages are indicated in the charts. Scale bars: number of mutations per site.

**Appendix 1**. HybPiper analysis. **(a)** Genome data from HybPiper analysis based on targeted gene capture in 132 Loliinae taxa. Taxon (name and authority), Sample code (Arbor Biosciences). Number and percentages of genes per taxon, sequence length and percentage of sequence length of genes (bp) per taxon. Read mapping efficiency parameters and assembly completeness at various thresholds (25%, 50%, 75%) of 353 genes. Sequence length of each of the 351 studied genes in the Loliinae taxa (numbers correspond to genes' codes). Mean values across the studied samples and total values are indicated at the end of the table. (**b**) Summary statistics and post-cleaning characteristics of the 234 single-copy multiple gene alignment using AMAS software.

Appendix 1 is available at Github (<https://github.com/Bioflora/LoliinaeGeneCapture>) and Dryad (<https://datadryad.org/stash/share/zMQ9ck5vFmQ83TFgtLlCT5jjJkA1N7JXG5m4OlETjkg>: Hybridization episodes in Loliinae: DOI:10.5061/dryad.n2z34tn5j).

**Appendix 2**. Summary statistics and characteristics of the Loliinae single-copy-gene (scg-strict data set) supermatrix alignment and the Loliinae plastome alignment using AMAS software. Appendix 2 is available at Dryad (<https://datadryad.org/stash/share/zMQ9ck5vFmQ83TFgtLlCT5jjJkA1N7JXG5m4OlETjkg>: Hybridization episodes in Loliinae: DOI:10.5061/dryad.n2z34tn5j).
